# Enhancing Gastrodin Production in Yarrowia lipolytica by Metabolic Engineering

**DOI:** 10.1101/2024.03.10.584284

**Authors:** Yuanqing Wu, Shuocheng Li, Baijian Sun, Jingyi Guo, Meiyi Zheng, Aitao Li

## Abstract

Gastrodin, 4-hydroxybenzyl alcohol-4-O-β-D-glucopyranoside, has been widely used in the treatment of neurogenic and cardiovascular diseases. Currently, gastrodin biosynthesis has been achieved in model microorganisms. However, the production levels are insufficient for industrial applications. In this study, we successfully engineered a Yarrowia lipolytica strain to overproduce gastrodin through metabolic engineering. Initially, the engineered strain expressing the heterologous gastrodin biosynthetic pathway, which comprises chorismate lyase, carboxylic acid reductase, phosphopantetheinyl transferase, endogenous alcohol dehydrogenases, and a UDP-glucose dependent glucosyltransferase, produced 1.05 g/L of gastrodin from glucose in a shaking flask. Then, the production was further enhanced to 6.68 g/L with a productivity of 2.23 g/L/day by over-expressing the key node DAHP synthases of the shikimate pathway and alleviating the native tryptophan and phenylalanine biosynthetic pathways. Finally, the best strain, Gd07, produced 13.22 g/L of gastrodin in a 5-L fermenter. This represents the highest reported production of gastrodin in an engineered microorganism to date, marking the first successful de novo production of gastrodin using Y. lipolytica.

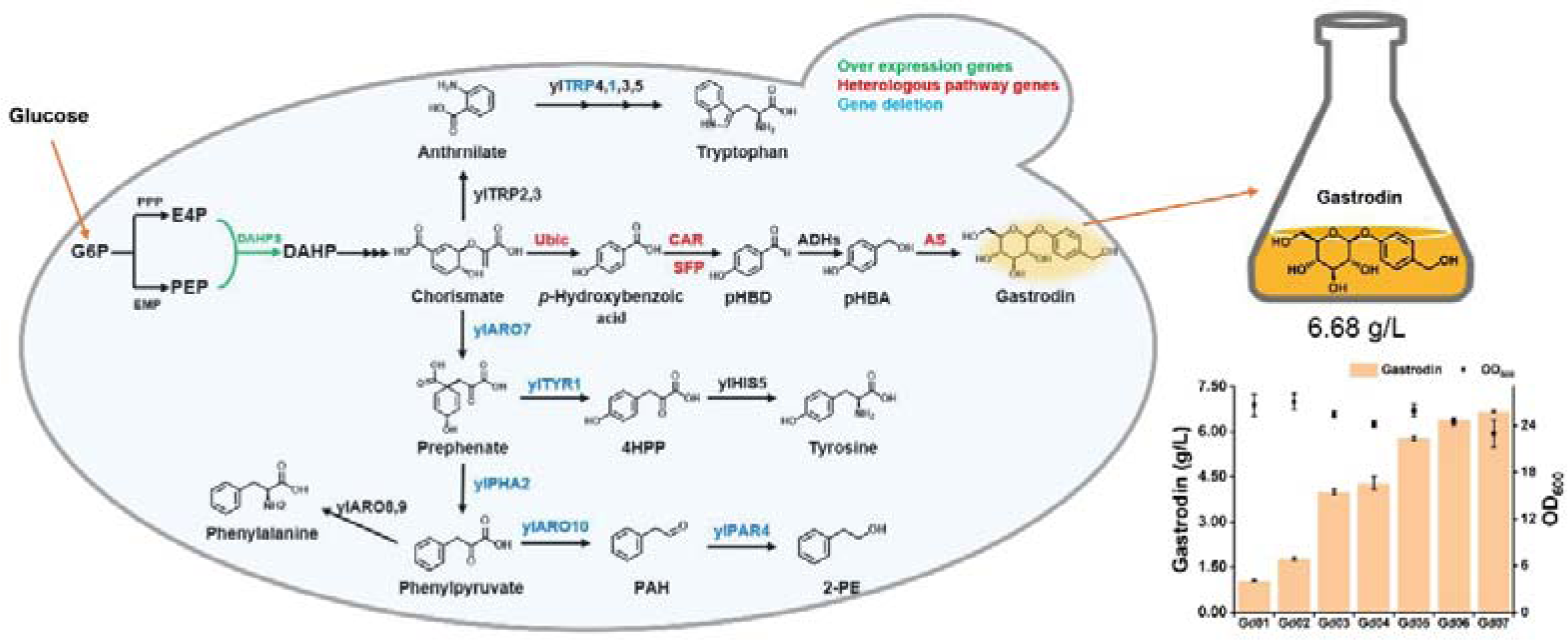

## INTRODUCTION

Gastrodin, a phenolic glycoside, is a major bioactive component in the traditional Chinese medicine *Gastrodia elata*. It is renowned for its pharmacological activities and has been widely used to treat central nervous system and cardiovascular symptoms ^1^. Researches indicate that gastrodin may protect against osteoporosis by reducing reactive oxygen species, protect human umbilical vein endothelial cells from homocysteine-induced injury, address intracerebral hemorrhage, mitigate steatohepatitis by activating the AMPK pathway, and facilitate blood vessel regeneration ^2–6^. As further research unveils additional bioactivities ^7^, the market demand for gastrodin is expected to grow.

Currently, gastrodin is primarily produced through the extraction of *G*. *elata,* but it suffers from several drawbacks including: 1) the natural habitat of *G*. *elata* is constrained by environmental and geographical factors since it is a saprophytic perennial herb; 2) the content of gastrodin varies depending on the growth conditions of *G*. *elata*, resulting in unstable yield and quality; 3) although artificial cultivation techniques for *G*. *elata* have been developed, the life cycle of the plant is lengthy ^8^. Chemical synthesis is another method for gastrodin production. However, the process requires complex reaction steps and large amounts of phosphorus and bromine, making it a low-yield, highly toxic, expensive, and environmentally unfriendly option ^9^. Therefore, there is an urgent need for a sustainable, efficient, and environment-friendly method to produce gastrodin.

Advances in biotechnology have led to the development of natural transformation systems involving fungi and plant tissue to produce gastrodin from *p*-hydroxybenzyl alcohol (pHBA) or *p*-hydroxybenzaldehyde (pHBD) ^9–11^. Although these methods have expanded gastrodin resources and increased production, they are time-consuming, labor-intensive, and expensive. In recent years, engineered heterologous microbial cells, including *Escherichia coli* (*E. coli*) and *Saccharomyces cerevisiae* (*S. cerevisiae*) have been used to produce gastrodin ^12–15^. Notably, the Liu group has established a novel microbial pathway for gastrodin synthesis from glucose by incorporating *Nocardia* carboxylic acid reductase (CAR), *Bacillus subtilis* phosphopantetheinyl transferase (SFP), endogenous alcohol dehydrogenases (ADHs), and *Rhodiola* glycosyltransferase UGT73B6. Finally, after genetic modification *via* metabolic and protein engineering, the engineered *E. coli* and *S. cerevisiae* strains produced 545 mg/L and 2.1 g/L of gastrodin, respectively, ^12, 15^. However, the production levels and productivities are considered insufficient for large-scale fermentation. The challenges in achieving high-level gastrodin production are attributed to the strong toxicity of intermediate aromatic compounds (such as flavanols and ferulic acid) to *E. coli* and *S. cerevisiae* cells because of their strong hydrophobicity and the weak hydrophobicity of the intracellular environment ^16^. Additionally, the intricate and diverse regulation within *E. coli* and *S. cerevisiae* cells limits the metabolic flux for aromatic synthesis, posing another obstacle ^17^. Therefore, it is necessary to develop a new microorganism cell factory for high-level gastrodin production.

*Yarrowia lipolytica,* a nonconventional oleaginous yeast, is generally regarded as safe (GRAS)^18^. In contrast to *S*. *cerevisiae*, *Y*. *lipolytica* possesses distinct physiological, genetic, and molecular biological characteristics, including a wide range of pH tolerance (2–9), high acetyl-CoA, pentose phosphate, tricarboxylic acid cycle (TCA), and lipid biosynthesis fluxes ^19–21^. Furthermore, *Y*. *lipolytica* secretes proteins in rich medium and organic acids in limited medium ^22^. Therefore, *Y*. *lipolytica* cells have been engineered to efficiently produce lipids, functional fatty acids, organic acids and tetraterpenoids ^23–27^. Recent studies have demonstrated that *Y*. *lipolytica* is a promising platform for the production of aromatic compounds. For example, Gu et al. ^28^ systematically modified the shikimate pathway and its precursor supply in *Y*. *lipolytica*, resulting in the highest production of 2-phenylethanol (2426.22 ± 48.33 mg/L), violacein (366.30 ± 28.99 mg/L), and deoxyviolacein (55.12 ± 2.81 mg/L) from glucose in shaking flask cultures. Additionally, the Borodina and Zhou groups combined feedback-inhibited relieving and precursor supply enhancement of the shikimate pathway with multicopy integration of heterologous genes, achieving resveratrol production of 12.4 g/L and 22.5 g/L in controlled fed-batch fermentor, respectively ^29, 30^. Notably, Shang et al. ^31^ constructed an efficient *Y*. *lipolytica* cell for arbutin production (glycosylation product of hydroquinone) from glucose through shikimate pathway carbon flux and fermentation optimization, with the highest production reaching 8.6 ± 0.7 g/L in shaking flask cultures. These findings suggest that *Y. lipolytica* is an excellent host for efficient gastrodin production.

In this study, we aimed to develop an efficient *Y*. *lipolytica* cell factory for gastrodin production. Strain Po1fΔ*Ku70* ^28^ was initially engineered to assess its synthetic capacity for gastrodin production. The heterologous pathway of gastrodin was first constructed through feeding experiments, resulting in the strain Gd01 producing 1.05 g/L of gastrodin from glucose in a shaking flask. Subsequently, overexpression of the shikimate pathway and alleviation of competitive pathways were performed to enhance gastrodin production. This led to the recombinant strain Gd07 achieving a remarkable milestone, producing 6.68 g/L of gastrodin in shaking flasks within 72 hours, with a productivity of 2.23 g/L/day. Finally, strain Gd07 produced 13.22 g/L of gastrodin in a 5-L fermenter. This represents the highest reported level of microbially produced gastrodin.

## RESULTS AND DISCUSSIONS

Gastrodin production from p-hydroxybenzoic acid in Y. lipolytica. Carboxylic acid reductase (CAR), phosphopantetheinyl transferase (SFP), endogenous alcohol dehydrogenases (ADHs), and glucosyltransferase constitute an artificial pathway that converts *p*-hydroxybenzyl alcohol (pHBA) into gastrodin (Figure. 1). The crucial step in this pathway is the conversion of *p*-hydroxybenzoic acid to *p*-hydroxybenzaldehyde (pHBD) catalyzed by CAR, which requires SFP post-translational activation and consumes ATP and NADPH. MAB4714, a CAR from *Mycobacterium abscessus* ATCC 19977, exhibits superior performance in the reduction of benzoic acid to benzaldehyde when co-expressed with SFP from *Bacillus subtilis in vitro* ^32^. Therefore, MAB4714 was used in the present study. *Y*. *lipolytica* demonstrates the ability to provide adequate ATP and NADPH by maintaining high fluxes of TCA and pentose phosphate pathway^24^. Furthermore, nine ADHs, one aldehyde ketone reductase (YALI0_B07117p) and one aldehyde dehydrogenase (YALI0_B01298p) have been identified and characterized in *Y*. *lipolytica* ^33, 34^, enabling the effective conversion of aromatic aldehydes into their corresponding alcohols. Thus, we speculated that the expression of CAR and SFP could efficiently convert *p*-hydroxybenzoic acid into pHBA. Then, cassette P_TEFin_-CAR-T_XPR2_-P_hp4d_-SFP-T_XPR2_ was integrated into a chromosome of *Y*. *lipolytica* strain Po1fΔ*Ku70 via* plasmid pINA1312, resulting in recombinant strain SC01. As anticipated, a new peak coincided with the retention time of the pHBA standard was observed during the HPLC analysis of the supernatant from strain SC01 after the addition of *p*-hydroxybenzoic acid (Figure 2A). This peak signified the effective transformation of the substrate, with a conversion exceeding 97% within 72 h of cultivation, leading to the production of *ca.* 1.10 g/L of pHBA (Figure 2B). Additionally, we observed an increase in the pHBD concentration to 0.21 g/L within the initial 48 h, followed by a decline to 0.01 g/L by the end of the experiment (Figure 2B). These results suggested that the co-expression of MAB4714 and SFP exhibited satisfactory performance in the strain Po1fΔ*Ku70*. Subsequently, we investigated the potential of an UDP-glucose-dependent glucosyltransferase from *Rauvolfia serpentina*, referred to arbutin synthase (AS), which has previously been employed for arbutin and gastrodin production in *E*. *coli* and *S*. *cerevisiae* ^15, 35^, to glycosylate pHBA in *Y*. *lipolytica*. The cassette P_hp4d_-AS-T_XPR2_ was integrated into the plasmid pINA1312 and transformed into strain Po1fΔ*Ku70*, resulting in the recombinant strain SC02. pHBA feeding experiments showed the presence of a new peak consistent with the gastrodin standard in the HPLC analysis of the cultivation broth of strain SC02 (Figure 2A). Consequently, strain SC02 produced 0.38 g/L of gastrodin from 1.24 g/L of pHBA at 72 h (Figure 2C). During the cultivation process, pHBA was also converted into pHBD (Figure 2C), which could be attributed to the enzymes (YALI0_B07117p or YALI0_B01298p) catalyzed reversible oxidative reactions ^33^.

**Figure 1.**
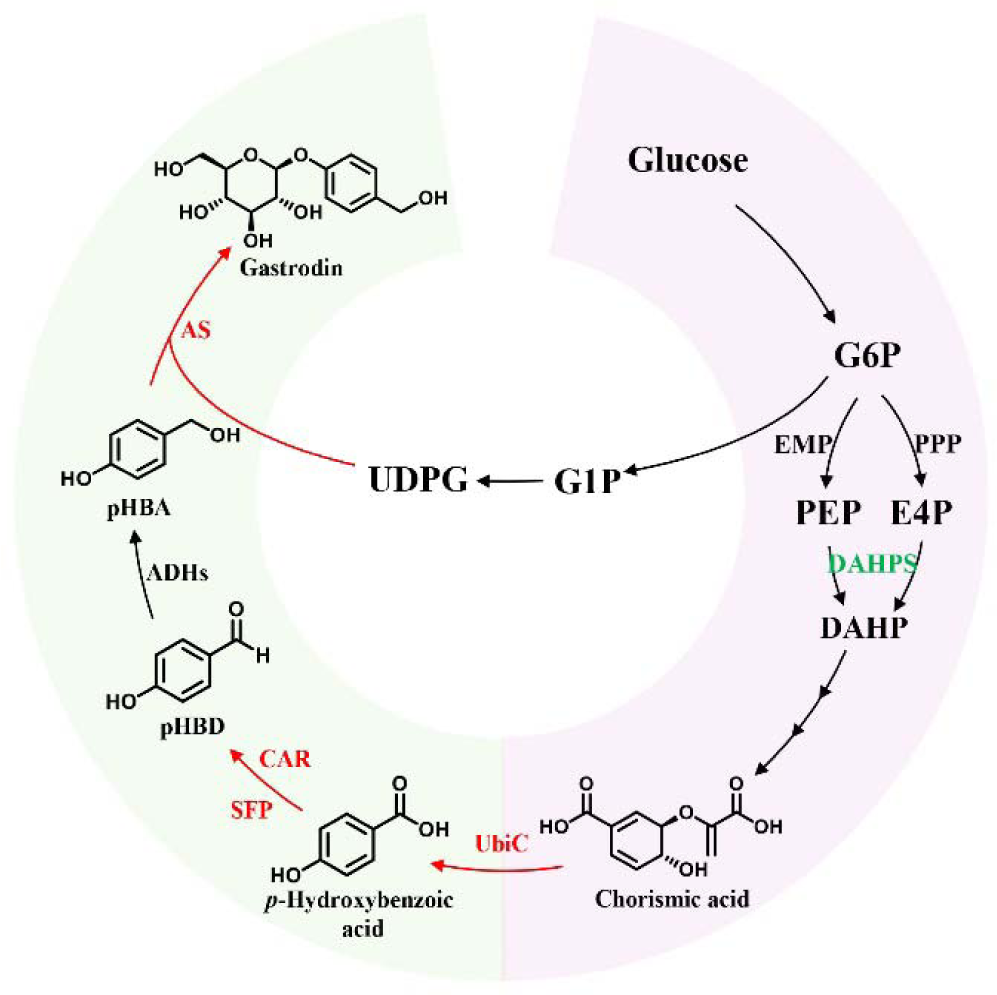
Engineered pathway for *the de novo* production of gastrodin from glucose in *Y*. *lipolytica*. G6P, 6-phosphate-D-glucose; G1P, 1-phosphate-D-glucose; UDPG, uridine diphosphate glucose; PEP, phosphoenolpyruvate; E4P, derythrose-4-phosphate; DAHP, 3-deoxy-D-arabino-heptulosonic acid 7-phosphate; pHBD, *p*-hydroxybenzaldehyde; pHBA, *p*-hydroxybenzyl alcohol; PPP, pentose phosphate pathway; EMP, glycolysis; DAHPS, 3-deoxy-D-arabino-heptulosonic acid 7-phosphate synthase; UbiC, chorismite lyase; CAR, carboxylic acid reductase; SFP, phosphopantetheinyl transferase; ADHs, alcohol dehydrogenases; AS, glycosyltransferase. Red arrows and enzymes represent the artificial pathway for gastrodin production.

**Figure 2.**
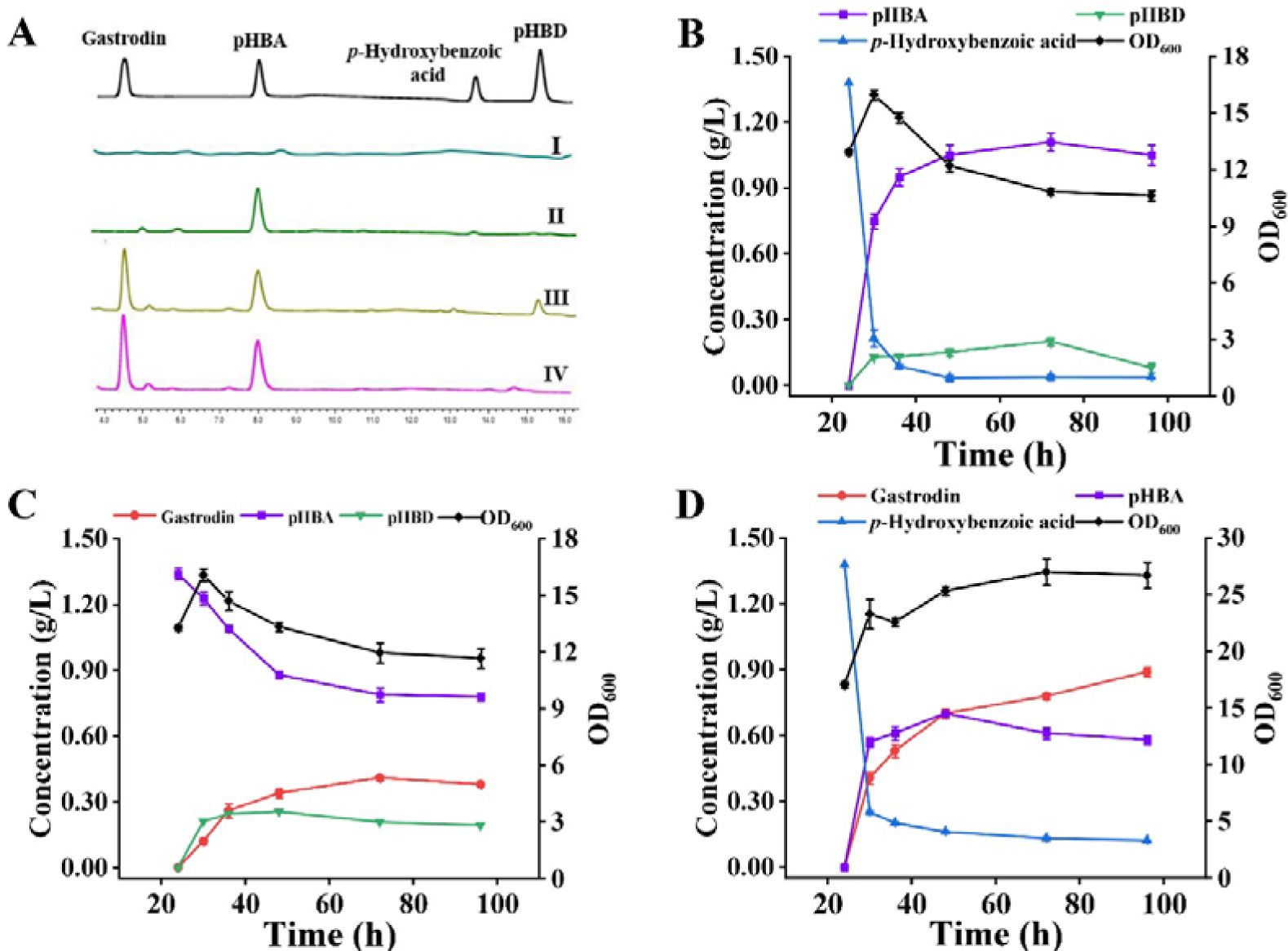
Product profiles of engineered *Y*. *lipolytica* strains through feeding experiments. (A) HPLC profiles of the supernatants produced by strains Po1fΔ*Ku70*(I), SC01(II), SC02 (III), and SC04 (IV). (B, C, and D) The product profiles produced by the recombinant strains SC01, SC02, and SC04. In each case, substrate concentrations of 1.38 g/L of *p*-hydroxybenzoic acid, 1.24 g/L of pHBA, and 1.38 g/L of *p*-hydroxybenzoic acid were utilized. Three biological replicates were performed for each strain, and the error bars represent standard deviations.

The above results demonstrated that the heterologous enzymes CAR and AS have high activities on the respective substrates in strain Po1fΔ*Ku70*. To produce gastrodin from *p*-hydroxybenzoic acid, the expression cassette P_TEFin_-CAR-T_XPR2_-P_hp4d_-SFP-T_XPR2_-P_hp4d_-AS-T_XPR2_ was inserted into the plasmid pINA1312 and transformed into strain Po1fΔ*Ku70* to create strain SC03. However, the strain SC03 produced a large amount of pHBA with formation of a trace amount of gastrodin when 1.38 g/L of *p*-hydroxybenzoic acid was added into the culture (Figure S2). This may be attributed to the impact of the integration site on heterologous gene expression within the genome of *Y*. lipolytica ^36^. Another possible reason is the low specificity of gene integration caused by the zeta region of plasmid pINA1312, resulting in unpredictable integration and unstable gene expression ^37^. Alternatively, the cassette P_hp4d_-AS-T_XPR2_ was inserted into the plasmid pINA1269 and transformed into strain SC01, resulting in strain SC04. Expectedly, strain SC04 produced 0.89 g/L of gastrodin at 72 h after 1.38 g/L of *p*-hydroxybenzoic acid was fed into the culture (Figure 2A and D). Thus, we succeeded in engineering *Y. lipolytica* to produce gastrodin from pHBA by adjusting its integration site.

De novo biosynthesis of gastrodin in Y. lipolytica. Chorismate is the point connecting gastrodin synthetic pathway and the native shikimate pathway (Figure 1) ^12, 15^. However, *Y*. *lipolytica* lacks the ability to synthesize *p*-hydroxybenzoic acid from chorismate due to the absence of chorismate lyase ^31, 38^. To overcome the defect, Shang et al. expressed chorismate lyase (UbiC) from *E*. *coli* in *Y*. *lipolytica* and engineered the strain to produce *p*-hydroxybenzoic acid for arbutin production ^31^. Considering the effect of the genome integration site on gastrodin production (SC04 and SC03), the *ubiC* gene from *E*. *coli* was inserted into plasmids pINA1269 or pINA1312-CAR-SFP, resulting in plasmids pINA1269-P_TEFin_-UbiC-T_XPR2_ or pINA1312-P_TEFin_-UbiC-T_XPR2_-P_TEFin_-CAR-T_XPR2_-P_hp4d_-SFP-T_XPR2_. Subsequently, the former plasmid was linearized and transformed into strain SC01, whereas the latter was linearized and transformed into strain Po1fΔ*Ku*70. This resulted in the generation of the strains SC05 and SC06. In shaking cultivation, it was observed that strain SC05 produced significantly more pHBA than strain SC06, with a production of 0.23 g/L compared to 0.08 g/L for SC06 (Figure 3A). Feeding experiments for gastrodin production led us to conclude that expressing CAR alone at a specific genomic site was more effective than co-expressing it with other genes at the same site in *Y. lipolytica*. Then, the *AS* gene was inserted into plasmid p1269-UbiC to construct cassette P_TEFin_-UbiC-T_XPR2_-P_hp4d_-AS-T_XPR2_. The cassette was transformed into strain SC01 to obtain the strain Gd01. As anticipate, a peak consistent with the gastrodin standard was identified in the fermentation broth of strain Gd01, confirming the *de novo* production of gastrodin from glucose (Figure S2). Finally, strain Gd01 produced 1.05 g/L at the end of fermentation (Figure 3B), with a slightly higher productivity compared to the engineered *S. cerevisiae* strain rGS-HBA (0.35 g/L/day *vs* 0.30 g/L/day) ^15^. Interestingly, the strain yielded pHBA at concentrations below 0.03 g/L during fermentation, with no pHBD detected (Figure 3B). These findings indicate the critical role of correctly assembling the heterologous pathway for natural product synthesis in *Y*. *lipolytica*. In addition, our results underscore *Y*. *lipolytica*’s significant potential as a highly hydrophobic host for the effective production of glycosylated compounds.

**Figure 3.**
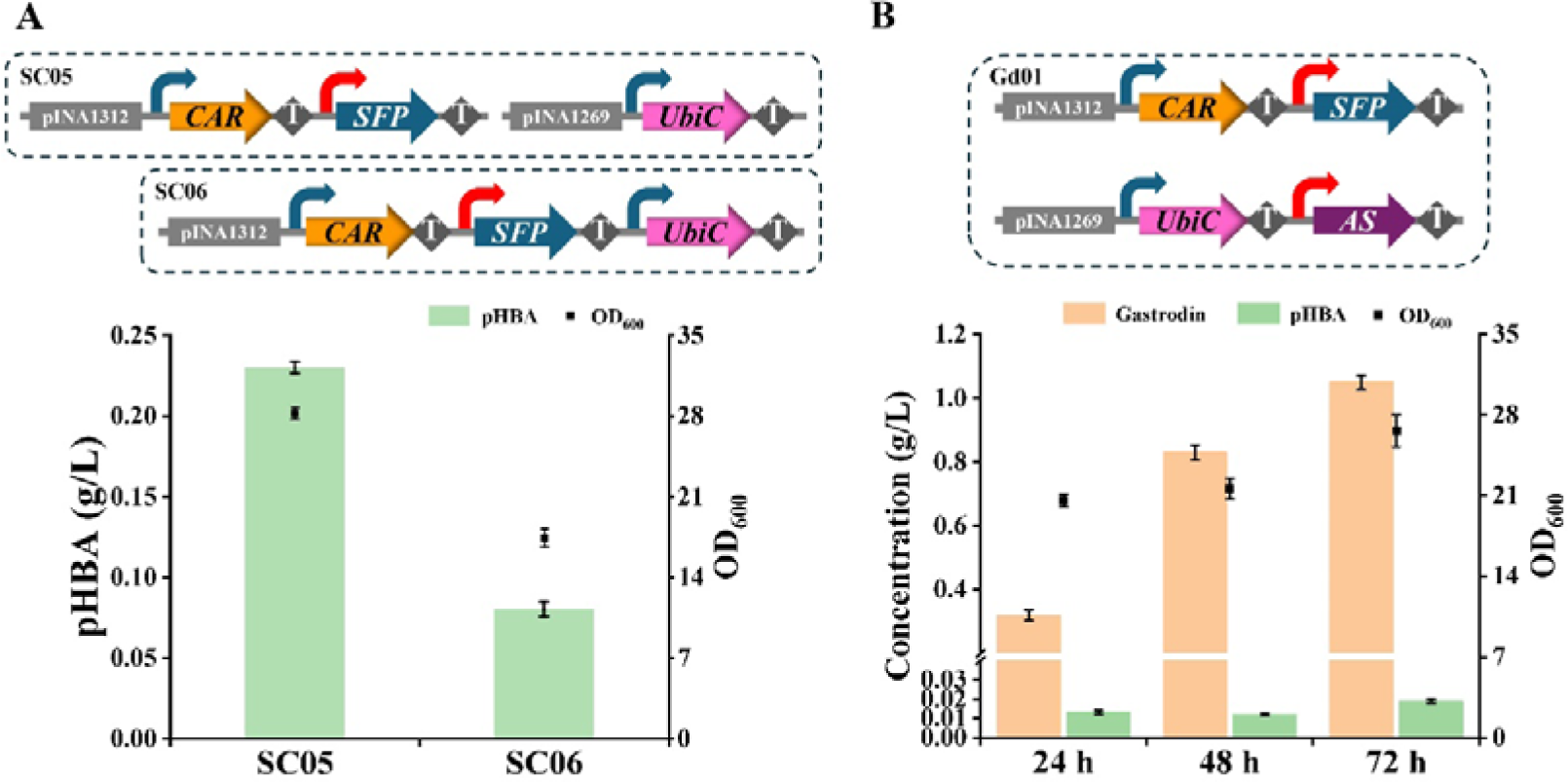
*De novo* biosynthesis of gastrodin in engineered *Y*. *lipolytica* strain Gd01. (A) pHBA production produced by strains SC05 and SC06, and (B) gastrodin production produced by strain Gd01 from glucose in YNB80 medium. Three biological replicates were performed for each strain, and the error bars represent the standard deviations. Blue and green curved arrows represent the promoters P_TEFin_ and P_hp4d_, respectively, and T represents the terminator T_XPR2_.

Overexpressing the bottleneck enzyme of the shikimate pathway to enhance gastrodin production. The condensation of phosphoenolpyruvate (PEP) and erythrose 4-phosphate (E4P), catalyzed by 3-deoxy-D-arabino-heptulosonate-7-phosphate synthase (DAHPS), is the initial and critical node of the shikimate pathway (Figure 1 and 4A). Previous studies have revealed the variability in enzymatic activity of DAHPS from different microorganisms, influenced by feedback regulation by aromatic amino acids ^17^. In *E. coli*, AroG contributes to 80% of the total DAHPS activity, and the feedback inhibition-insensitive mutant AroG^G146N^ is usually overexpressed to enhance the production of the desired products ^39, 40^. Similarly, in *Y*. *lipolytica*, overexpression of the isoenzymes DHS1 and DHS2 or feedback-insensitive version ylARO4 ^K221L^ has been demonstrated to significantly improve the production of aromatic compounds^28, 29, 31^. Thus, to enhance gastrodin production, the genes *DHS1*, *DHS2*, *ylARO4^K^*^221^*^L^*, and *aroG^G^*^146^*^N^*were targeted for overexpression.

**Figure 4.**
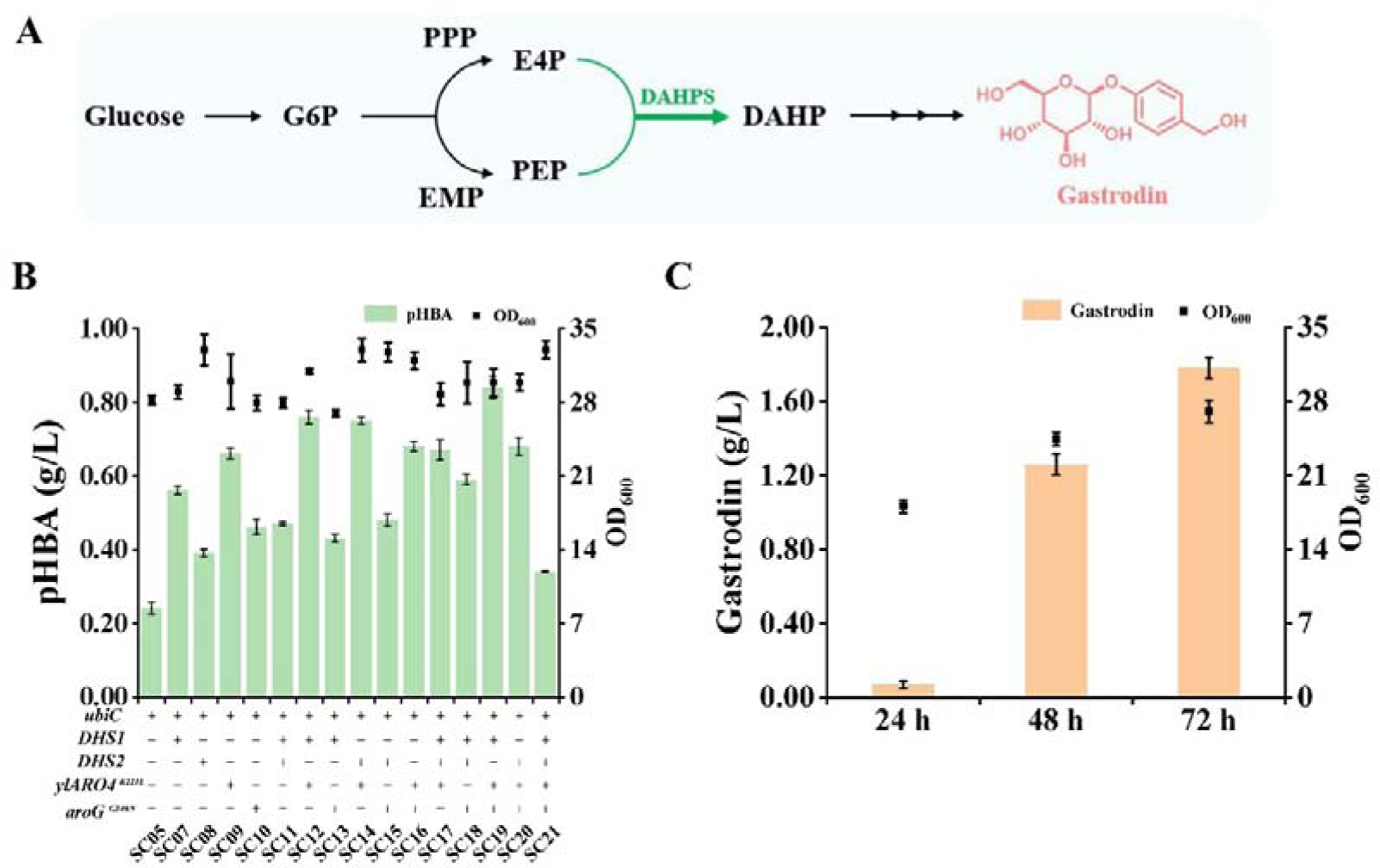
Gastrodin production in engineered *Y*. *lipolytica* strain Gd02 by strengthening DAHPS. (A) Production of pHBA by strains overexpressing DAHPS. (B) Gastrodin production produced by strain Gd02. Three biological replicates were performed for each strain, and the error bars represent standard deviations.

To obtain then strain with the highest gastrodin production, our initial focus was on assessing the impact of genes overexpression on the precursor production, pHBA. To achieve this goal, constructs were designed for cassettes P_hp4d_-DHS1-T_XPR2_, P_hp4d_-DHS2-T_XPR2_, P_hp4d_-ylARO4^K221L^-T_XPR2_, and P_hp4d_-aroG^G146N^-T_XPR2_, which were subsequently integrated individually or in combination into the plasmid pINA1269-UbiC. This resulted in the generation of 15 new expression cassettes. Then, the cassettes were integrated into the genome of strain SC01, resulting in strains SC07–SC21. Cultivation in shaking flask revealed that overexpression led to increased pHBA production compared to strain SC05, with improvements ranging from 41.9% to 256.4% at 72 h (Figure 4A). Notably, strains SC12, SC14, and SC19 yielded 0.76 g/L, 0.75 g/L, and 0.84 g/L of pHBA, representing increases of 220.8%, 219.1%, and 256.4%, respectively. We observed that strain SC21 exhibited a modest 41.9% improvement in pHBA production, likely due to the significant metabolic burden caused by the concurrent expression of four enzyme genes ^41, 42^. Thereafter, the cassette P_hp4d_-AS-T_XPR2_ was introduced into strain SC19 *via* plasmid pINA1269, resulting in the gastrodin-producing strain Gd02. As expected, the strain produced 1.78 g/L of gastrodin after fermentation for 72 h (Figure 4B), which was 69.5% higher than that produced by strain Gd01. These results effectively demonstrated that reinforcing the condensation reaction of E4P and PEP can significantly boost the production of targeted chemicals in *Y*. *lipolytica*.

Alleviating competitive pathways to further enhance gastrodin production. The biosynthesis of aromatic amino acids is a competitive pathway for gastrodin production, and knockout of these pathways may increase gastrodin production (Figure 5A). Gao et al. ^43^ demonstrated that the deletion of *ylTRP1* gene caused tryptophan autotrophy in *Y*. *lipolytica*, indicating that deletion of the *TRP1* gene effectively hinders the carbon flux towards tryptophan biosynthesis. The *ylARO7* gene, encoding chorismate mutase, is the crucial node of L-tyrosine and L-phenylalanine (Phe) biosynthesis, and its overexpression promoted the production of targeted products ^28, 29^. Thus, genes *ylTRP1* and *ylARO7* were respectively deleted in strain SC19 *via* the CRISPR/Cas9 system, generating strains SC22 (SC19, *ylTRP1*^-^), SC23 (SC19, *ylARO7*^-^), and SC24 (SC19, *ylTRP1*^-^, *ylARO7*^-^). The three strains were cultivated in a shaking flask for 72 h to assess the production of pHBA. As shown in Figure 5B, it is evident that strain SC22 yielded 1.95 g/L pHBA, marking a 131.9% increase in comparison to strain SC19. Conversely, strains SC23 and SC24 only yielded 0.12 g/L and 0.11 g/L, indicating a decrease of 85.5% and 86.6% respectively. Moreover, the OD_600_ value of strain SC22 notably decreased by 67.4%, whereas strains SC23 and SC24 showed minimal changes compared to their parent strain. These observations might be related to the altered distribution of chorismite carbon flow caused by the deletion of genes *ylTRP1* and *ylARO7*, which needs to be explored to understand the underlying mechanisms through ^13^C metabolic flux analysis in the future work ^44^. Then, the *AS* gene was integrated into the genome of strain SC22, resulting in strain Gd03. However, the productions of gastrodin and biomass produced by the strain were significantly decreased in YNB80 medium after fermentation for 72 h in a shaking flask (Figure 5C). We speculated that gene *ylTRP1* deletion affected the amino acid metabolism of the strains, thereby decreasing their biomass. Subsequently, different concentrations of tryptophan and tryptone were added to YNB80 for the cultivation of strain Gd03, respectively. Unexpectedly, tryptophan supplementation had no effect on gastrodin (Figure S3), whereas tryptone supplementation greatly increased gastrodin production (Figure 5C). Finally, strain Gd03 achieved a gastrodin production of 4.01 g/L upon the supplementation of 8 g/L of tryptone, marking a 124.9% improvement over strain Gd02, while cell growth resumed to its usual levels. Additionally, deletion of the gene *ylTYR1* in strain Gd03 did not enhance gastrodin production.

**Figure 5.**
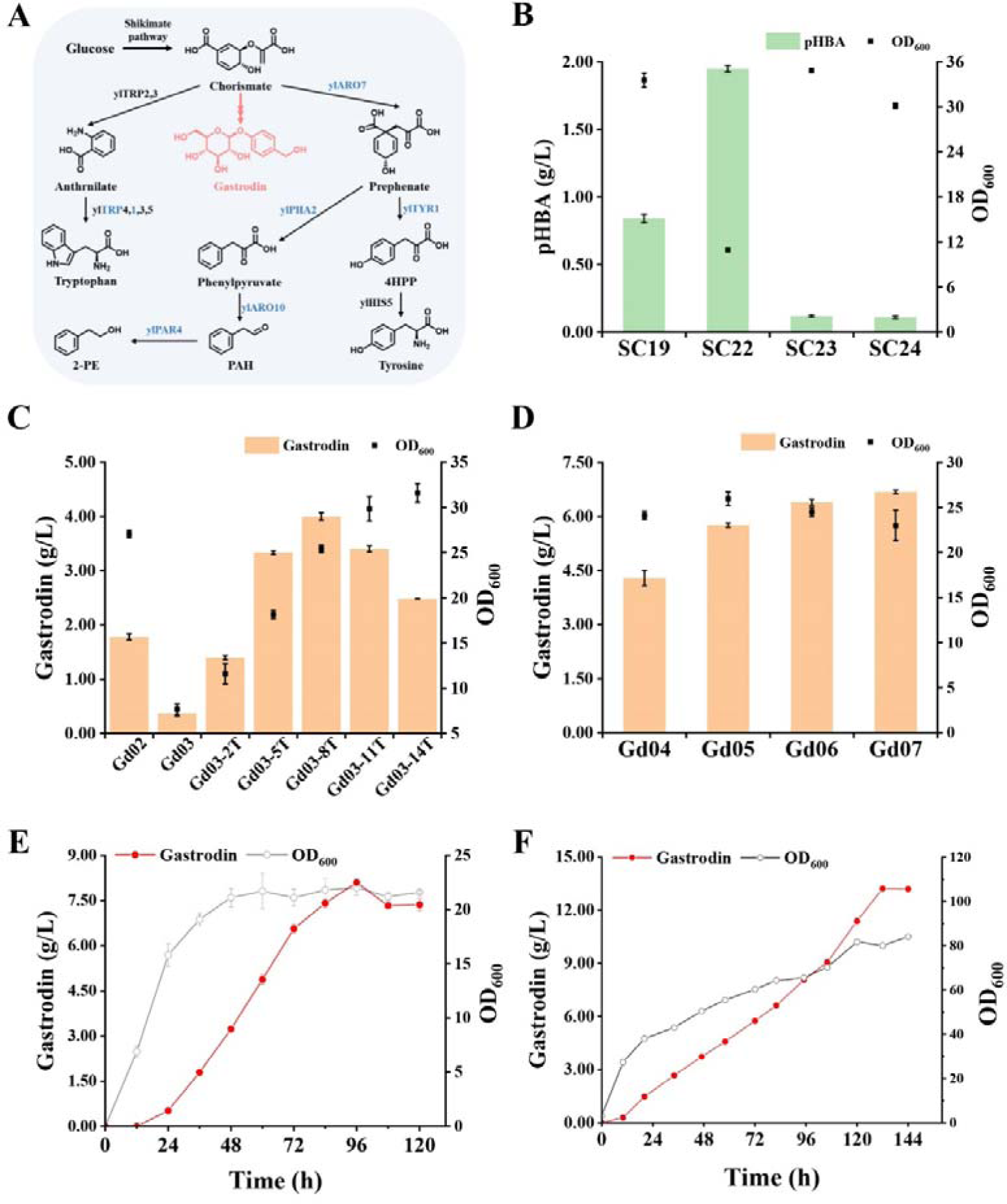
Alleviating competitive pathways to increase gastrodin production. (A) Endogenous competitive pathways for gastrodin production. ylTRP2, anthranilate synthase; ylTRP3, anthranilate synthase; ylTRP4, anthranilate phosphoribosyltransferase; ylTRP1, phosphoribosylanthranilate isomerase; ylTRP5, tryptophan synthase; ylARO7, chorismate mutase; ylTYR1, prephenate dehydrogenase; ylPHA2, prephenate dehydratase; ylARO10, phenylpyruvate decarboxylase; ylPAR4, phenylacetaldehyde reductases. 4HPP, 4-hydroxyphenylpyruvate; PAH, phenylacetaldehyde; 2-PE, 2-phenylethanol. Red indicates the heterologous pathway, and blue represents deletion. (B) The effects of deleting genes *ylTRP1* and *ylARO7* on pHBA production in strain SC19. (C) The effect of tryptone supplementation on gastrodin and biomass production in strain Gd03. 2T, 5T, 8T, 11T, and 14T represent 2, 5, 8, 11, and 14 g/L of tryptone, respectively. (D) The impact of attenuating 2-PE carbon flux on gastrodin production based on strain Gd03. The strains were assessed in YNB80 medium containing 8 g/L tryptone. (E) Gastrodin production produced by strain Gd07 after extended shaking flask cultivation. (F) Fed-batch fermentation of strain Gd07 in a 5-L fermenter.

Subsequently, strain Gd03 was found to produce 0.23 g/L of 2-phenylethanol (2-PE), while the deletion of gene *ylARO7* was unfavourable for increasing pHBA flux. In an effort to further improve gastrodin production, the flux through the Ehrlich pathway, which is known for efficient 2-PE production in *Y*. *lipolytica* ^34, 45^, was attenuated. The three pathway genes *ylARO10* encoding phenylpyruvate decarboxylase, *ylPAR4* encoding phenylacetaldehyde reductase, and *ylPHA2* encoding prephenate dehydratase were successively deleted in strain Gd03, resulting in strains Gd04 (Gd33, *ylARO10*^-^), Gd05 (Gd33, *ylARO10*^-^, *ylPAR4*^-^), and Gd06 (Gd33, *ylARO10*^-^, *ylPAR4*^-^, *ylPHA*2^-^), respectively. The three strains were assessed for gastrodin production in YNB80 medium supplemented with 8 g/L tryptone. As a result, the gastrodin production of the three strains reached 4.29 g/L, 5.76 g/L, and 6.39 g/L, representing increases of 4.0%, 39.6%, and 54.8% compared to the starting strain Gd03, respectively (Figure 5D). Accordingly, pHBA production decreased by 17.4%, 40.8%, and 64.5%, while 2-PE production slightly decreased (Figure S3). The removal of genes ylARO10 and ylPAR4 significantly boosted Phe production by 166.4%, whereas the deletion of gene ylPHA2 led to a notable decrease of 23.2% compared to that of strain Gd03 (Figure S5). To attempt to decrease the production of 2-PE, we identified another phenylpyruvate decarboxylase encoding gene *YALI0D10131g via* NCBI blast. Therefore, this gene was deleted in strain Gd06, generating strain Gd07. Results showed that the production of gastrodin produced by the strain reached 6.68 g/L after 72 h of fermentation, with only a 4.5% increase compared to strain Gd06 (Figure 5D). The production and productivity were 3.2- and 7.4-fold higher than those of engineered *S*. *cerevisiae* strain rGS-HBA (6.68 g/L *vs* 2.1 g/L, and 2.23 g/L/day *vs* 0.3 g/L/day) ^15^. The productions of pHBA and 2-PE produced by Gd07 was not affected, but Phe production decreased by 13.6% compared to strain Gd06 (Figure S5). Although we failed to significantly reduce 2-PE production, our findings indicate that decreasing carbon flux for 2-PE production impacted the levels of pHBA and Phe production, resulting in an increase in gastrodin production in *Y*. *lipolytica*. The next crucial step involves the investigating the underlying mechanism of this phenomenon through multi-omics approaches and making adjustments to further enhance gastrodin production using CRISPRi or CRISPRa systems ^46, 47^. Furthermore, although the production of 2-PE generated by these modified strains was not substantial, additional genes could be targeted for deletion to further enhance gastrodin production in the future ^34^. Subsequently, the highest gastrodin production produced by strain Gd07 reached 8.10 g/L when the cultivation time was extended to 120 h in shaking flask (Figure 5E). Finally, strain Gd07 was fed-batch cultivated in a 5-L fermenter, the gastrodin production reached 13.22 g/L at 132 h (Figure 5F), marking the highest reported yield for the engineered strains up to date. Undoubtedly, the production via fed-batch fermentation is unsatisfactory. The main reason is likely the improper C/N ratio resulting from the deletion of the gene *ylTRP1*, which significantly reduces the biomass as well as the production of pHBA and gastrodin (Figure 5B and C). Another contributing factor is the inappropriate concentration of residual glucose in the fermenter because one UDPG derived from glucose is required to produce one gastrodin (Figure 1). In the future, improving fermentation efficiency will involve optimizing the fermentation process, including medium and fed-batch optimization, through expert operation and artificial intelligence strategies^48, 49^.

## CONCLUSIONS

In conclusion, a successful gastrodin production was achieved in *Y*. *lipolytica* by integrating a heterologous pathway consisting of UbiC, CAR, SFP, and AS enzymes into the yeast genome using a combination of genomic integration sites. By enhancing the carbon flux of the shikimate pathway and reducing the carbon flux of competitive pathways, we achieved the highest reported gastrodin production and productivity to date. In contrast to engineered model microorganisms that produce gastrodin, our work resulted in the construction of an efficient non-conventional yeast *Y. lipolytica* cell factory for gastrodin production, offering an over-producing strain suitable for industrial-scale production, thus presenting a promising alternative to existing production methods.

## METHODS

Plasmids, Strains, and Medium. The plasmids and strains used in this study are listed in Table 1 and S1. *E*. *coli* DH5α was used to construct *Y*. *lipolytica* recombinant plasmids. *E*. *coli* cells were cultured at 37 °C in solid Luria-Bertani (LB) medium (5 g/L yeast extract, 10 g/L tryptone, and 10 g/L NaCl) supplemented with appropriate antibiotics at a final concentration of 100 μg/mL ampicillin or 50 μg/mL kanamycin. The *Y*. *lipolytica* strain Po1fΔ*Ku70* with non-homologous end-joining reduction, leucine, and uracil auxotrophy was used as the starting strain for the construction of all engineered strains. The supplemental yeast nitrogen base (YNB) medium (6.7 g/L yeast nitrogen base, 10 g/L glucose, and 15 g/L Bacto agar without leucine and uracil) was used to screen transformants. *Y*. *lipolytica* strains were cultured in the complex medium yeast peptone dextrose (YPD) (20 g/L tryptone, 10 g/L yeast extract, and 20 g/L glucose). YNB80 medium (8 g/L yeast nitrogen base, 9 g/L yeast extract, and 80 g/L glucose) was used for feeding experiments and shake flask fermentation.

**Table 1.**
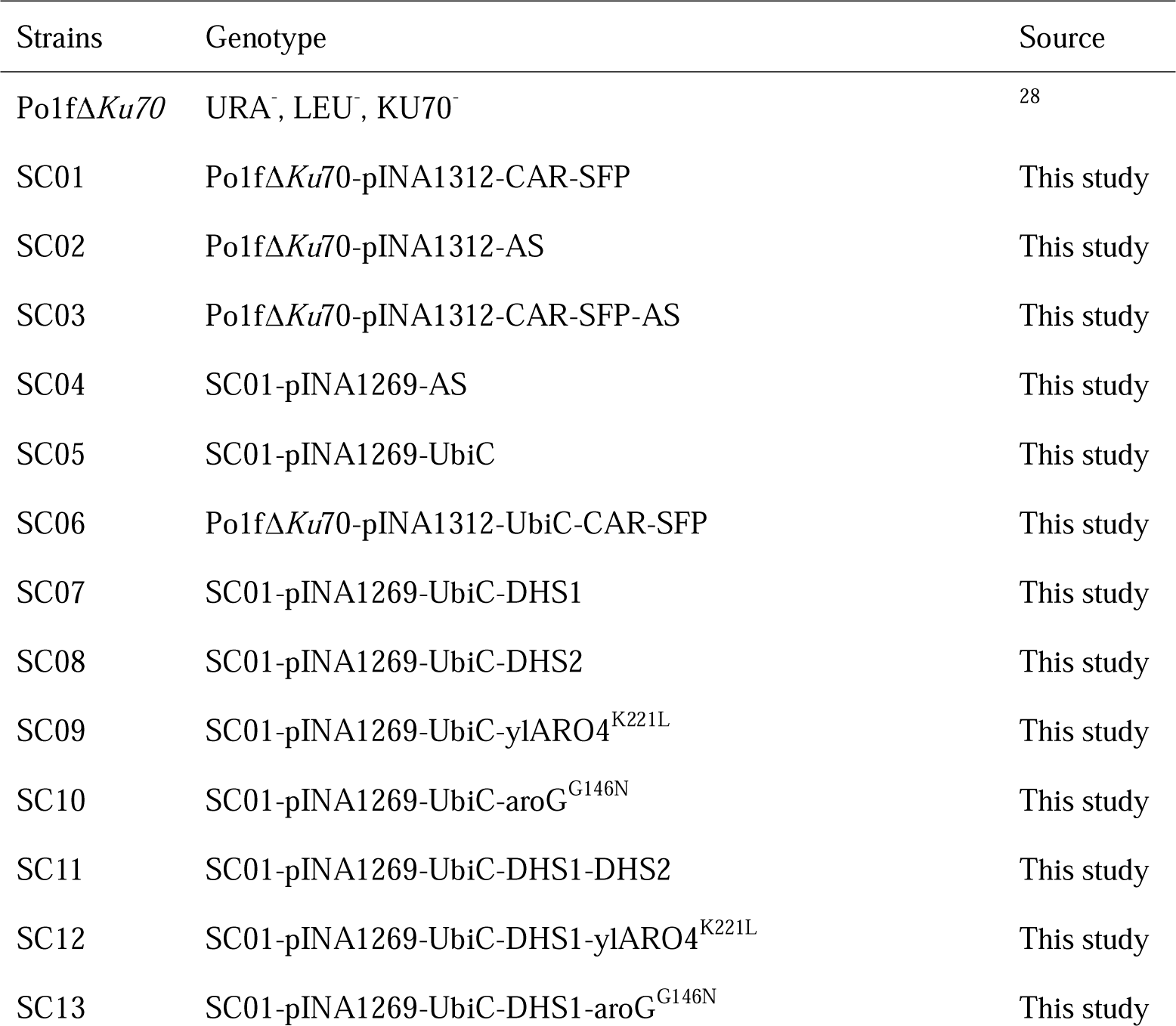
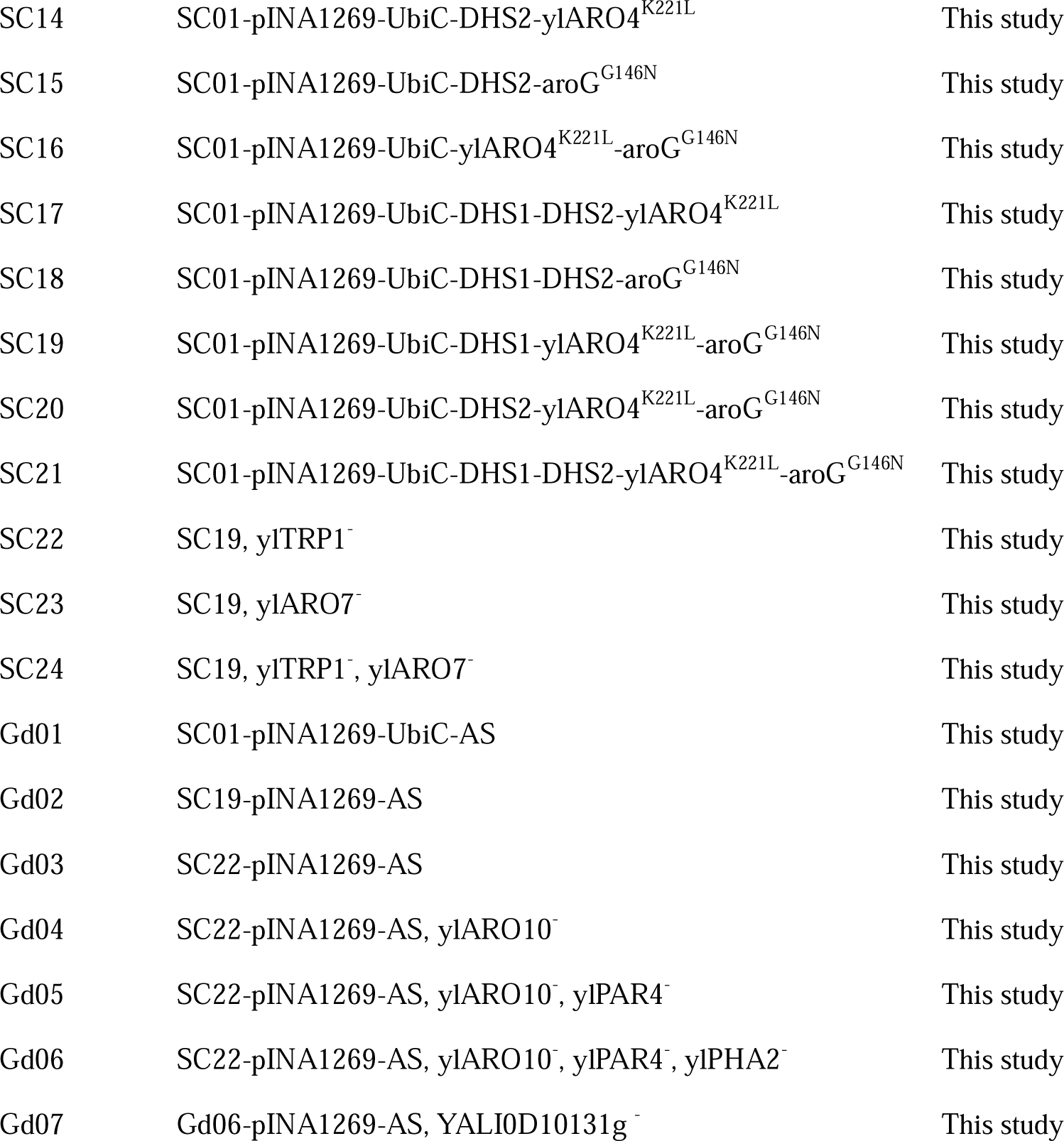
List of the Strains Used in This Study.

DNA Manipulation. The primers used for the amplification of the DNA fragments are listed in Table S2. The exogenous genes used for gastrodin synthesis, CAR (CAM64782.1), SFP (P39135.2), and AS (AJ310148.1), were codon-optimized and chemically synthesized by the Wuhan GeneCreate Biological Engineering Co. LTD (China). Polymerase chain reaction (PCR) was performed using Primer STAR Max DNA Polymerase (Takara) to amplify the target gene. The DNA fragment was purified using a purification kit (Omega Bio-tek Inc. Geogia, USA) and inserted into the plasmids pINA1312 or pINA1269 ^50, 51^, to construct an expression cassette via T5 exonuclease-dependent assembly ^52^. Genes are controlled by the hp4d or TEFin promoter and are terminated by the XPR2 terminator. For the expression of multiple genes, cassette DNA fragments were ligated into one plasmid using the Uniclone One Step Seamless Cloning Kit (Genesand, Beijing, China). DNA fragments of the TEFin promoter and shikimate pathway genes were amplified from the genome of *Y*. *lipolytica*, and the aroG gene was amplified from the genome of *E*. *coli*. Promoter hp4d and terminator XPR2 were from plasmid pINA1312 or pINA1269. All plasmid constructs were verified using PCR and sequenced (GeneCreate, Wuhan, China).

For gene knockout/knock-in, the CRISPR-Cas system was reconstructed. First, the DNA fragments of Cas9 from *Streptococcus pyogenes* controlled by the promoter TEFin and sgRNA-HDV controlled by the promoter SCR1′-tRNA^Gly^ ^43, 53^ were codon-optimized and chemically synthesized. Then, DNA fragments of Cas9, sgRNA-HDV with promoter SCR1′-tRNA^Gly^, TEFin promoter, and PEX20 terminator were amplified using the corresponding primers listed in Table S2. After purification using a kit, the four DNA fragments were integrated into the plasmid pRRQ2 using the Uniclone One Step Seamless Cloning Kit (Genesand, Beijing, China). Consequently, the recombinant plasmid pYlCas9 was created, and only the gRNA sequence needs to be replaced to build a new system. For target genes, sgRNA was designed using CHOPCHOP (http://chopchop.cbu.uib.no) and its secondary structure was checked using RNAfold WebServer (http://rna.tbi.univie.ac.at/cgi-bin/RNAfold.cgi). For gene knockout, the upstream homologous arm and the downstream arm of the targeted gene were amplified and ligated via overlapping PCR. For gene knock-in, DNA fragments of upstream homologous arms, promoters, targeted genes, terminators, and downstream arms were amplified and fused by overlapping PCR. The length of each arm is 1 kb.

Yeast transformation and cultivation. For gene overexpression mediated by plasmids pINA1312 or pINA1269, the constructed and corrected recombinant plasmids were digested with *Not* I for linearization and then transformed into Po1fΔ*Ku70* using the Frozen-EZ yeast transformation II Kit (Zymo Research). Transformants were screened on YNB plates supplemented with auxotrophic supplements containing appropriate amino acids and placed in an incubator at 30 °C for 2-3 day. A single colony was picked and verified using colony PCR and genomic DNA PCR ^31, 43^, and sequenced using sequencing primers to verify the successful integration of the target genes. For gene knockout/knock-in mediated by the CRISPR-Cas system, the constructed donor DNA fragment and recombinant plasmid pYlCas9 were simultaneously transformed into the targeted strain using the Frozen-EZ yeast transformation II Kit (Zymo Research). The amounts of donor DNA and plasmid pYlCas9 used were 1 μg and 500 ng, respectively. The transformants were screened by placing YNB plates without Leu and incubating them at 30 °C for 2-3 days. A single colony was picked and verified using colony PCR and genomic DNA PCR, and sequenced using sequencing primers to verify the successful integration of the target genes. To cure the plasmid pYlCas9, the correct colony harboring pYlCas9 was inoculated into 2 mL of YPD medium. The culture was incubated overnight, diluted, and spread onto YPD plates. The colonies were picked and transferred to fresh YPD and YNB plates without Leu and cultured under the same conditions for 2-3 days. Colonies curing pYlCas9 can grow on YPD plates but not on YNB plates without Leu. The colony cured with pYlCas9 was used for the next round of genome editing.

Feeding experiments. To evaluate the capacity of the substrates to be converted into products by recombinant *Y. lipolytica* strains, over 10 positive colonies were transferred from YNB plates with auxotrophic supplements (Leu or Ura) into 30-mL glass tubes (15 × 150 mm) containing 3 mL YPD medium and then incubated for 24 h at 30 L and 250 rpm. Subsequently, 500 µL of the culture was transferred into 50 mL of YNB80 medium in a 250-mL flask and cultivated at 30 °C and 250 rpm. After 24 h of incubation, certain amounts of substrate were added to the culture solutions and cultivated for 72 h. The samples were collected every 24 h after the substrates were added. Cell growth was measured by monitoring cell density at OD_600_, and production of the targeted chemicals was measured using high-performance liquid chromatography (HPLC). The optimal strain was chosen and preserved for the subsequent operation.

Shaking Fasks and Fed-Batch Fermentation. For each round of genetic modification in shaking flask fermentation, over 10 positive colonies were transferred from YNB plates with auxotrophic supplements (Leu or Ura) into 15-mL glass tubes (15 × 130 mm) containing 3 mL YPD medium and then incubated for 24 h at 30 L and 250 rpm. Samples were collected every 24 h to measure the OD_600_, glucose in the medium, and products. The optimal strain was chosen and preserved for the subsequent operation.

For fed-batch fermentation, the targeted strain culture was streaked onto an YPD plate and grown at 30 °C for 48 h. A single colony from the plate was transferred to 2 mL YPD in 15-mL glass tubes (15 × 130 mm) and incubated at 30 °C for 24 h. Then, 500 µL of the culture was transferred into 50 mL of YPD medium in a 250-mL flask and cultivated at 30 °C and 250 rpm for 24 h. The cells were obtained by centrifugation for 5 min at 5000 × *g*, washed twice with sterile water, and resuspended in 5 mL volume. The cell suspension was transferred to a 5-L fermenter with 2-L of fermentation medium, and the initial OD_600_ was 1.0. The fermenter was (Biotech-5JGY, Baoxing, Shanghai, China) equipped with sensing electrodes for pH, dissolved oxygen (DO), and temperature. One liter of medium contained 40 g D-glucose, 8 g tryptone, 9 g yeast extract, 3 g KH_2_PO_4_, 5 g (NH_4_)_2_SO_4_, 0.5 g MgSO_4_⋅ 7H_2_O, 2 mL trace metal solution, and 1 mL vitamin solution. The components of the trace metal and vitamin solutions were prepared as previous report ^54^. Fermentation was carried out at 30 °C with an airflow rate of 1.0 vvm, and DO was maintained at 30% by adjusting the agitation speed from 540 rpm to 1,000 rpm. The feed medium contained 500 g/L glucose, 50 g/L tryptone, a 10 × trace metal solution, and a 10 × vitamin solution. Samples were collected to determine OD_600_, glucose concentration, and product concentration.

Analytic methods. The OD_600_ and glucose in the broth were analyzed using an ultraviolet spectrophotometer (Beijing Puxi Universal, Co., Ltd, Beijing, China) and SBA-40C biosensor analyzer (Institute of Microbiology, Shandong, China). To quantify the production of the chemicals, 0.1 mL of fermentation broth and 0.9 mL of methanol was mixed thoroughly. The extract was then centrifuged at 12,000 × g for 10 min and filtered using a 0.22 μm filter membrane. The supernatants were analyzed using HPLC on a Shimadzu system equipped with a UV detector. For gastrodin, pHBA, pHBD and *p*-hydroxybenzoic acid, the column used was a Shim-pack GIST-HP C18, 4.6 × 150 mm, with a particle size of 5 μm. A low-pressure gradient methanol–water (containing 0.1% trifluoroacetic acid) system at a flow rate of 1.2 mL/min was used to separate the compounds. The HPLC program was as follows: 15% methanol for 6 min, 15–25% methanol for 4 min, 25% methanol for 3 min, 25–15% methanol for 2 min, and 25% methanol for 3 min. The products were detected at 220 nm and 40 °C. For 2-PE, analysis was carried out at 215 nm at 40 °C (column oven temperature) with a mobile phase containing 50% (v/v) methanol in water with 0.1% (*v*/*v*) formic acid at a flow rate of 1.0 mL/min equipped with a ZORBAX SB-C18 column (4.6 × 250 mm, 5 μm, Agilent). Quantification was performed using standard calibration curves prepared using a series of known concentrations of the standard. The R^2^ value of the standard curve is higher than 0.999. The titers are presented as mean ± standard deviation (SD).

## ASSOCIATED CONTENT

### Supporting Information

Plasmids (Table S1) and primers (Table S2) used in this study, the supernatant from the recombinant strains SC03, Gd01and Gd07 (II) were subjected to HPLC analysis (Figure S1), the effects of different tryptophan concentrations on gastrodin and biomass production in strain Gd02 (Figure S2), the productions of pHBA and 2-PE produced by different strains (Figure S3)

### Author Contributions

A. L. and Y.W. designed the study. S. L., B. S. and J. G. performed plasmid and strain construction, and fermentation experiments. A. L. and Y. W. analyzed all data. Y. W. wrote the manuscript. Y. W., S. L. and J. G. drew the figures. A. L. supervised the project and revised the manuscript. All authors have read and approved the final manuscript.

### Notes

The authors declare no competing financial interests.

## ACKNOWLEDGMENTS

This study was funded by the National Key Research and Development Program of China (2019YFA0905000), the Key Science and Technology Innovation Project of Hubei Province (2021BAD001), the Innovation Base for Introducing Talents of Discipline of Hubei Province (2019BJH021), and the Research Program of State Key Laboratory of Biocatalysis and Enzyme Engineering. We sincerely thank Dr. Yang Gu (School of Food Science and Pharmaceutical Engineering, Nanjing Normal University) for providing strain *Po1f*Δ*Ku70*.

