## Supplementary material for "Enhancing Gastrodin Production in Yarrowia lipolytica by Metabolic Engineering": Table S1, Table S2, Figure S1, Figure S2, Figure S3, Figure S4, Figure S5

**Table S1.** Plasmids used in this study

| Plasmid | | Description | Source |
| --- | --- | --- | --- |
|  | pINA1312 | *Y*. *lipolytica*-integrative plasmid, hp4d promoter, XPR2 terminator, ura3d1 selection marker, Km^R^ | ^1^ |
|  | pINA1269 | *Y*. *lipolytica*-integrative plasmid, hp4d promoter, XPR2 terminator, LEU2 selection marker, Amp^R^ | ^1^ |
| 1 | pINA1312-AS | pINA1312 plasmid harbors *AS* gene | This study |
| 2 | pINA1269-AS | pINA1269 plasmid harbors *AS* gene | This study |
| 3 | pINA1312-CAR | pINA1312 plasmid harbors *CAR* gene | This study |
| 4 | pINA1312-SFP | pINA1312 plasmid harbors *SFP* gene | This study |
| 5 | pINA1269-UbiC | pINA1269 plasmid harbors *ubiC* gene | This study |
| 6 | pINA1269-DHS1 | pINA1269 plasmid harbors *DHS1* gene | This study |
| 7 | pINA1269-DHS2 | pINA1269 plasmid harbors *DHS2* gene | This study |
| 8 | pINA1269- ARO4 ^K221L^ | pINA1269 plasmid harbors *ARO4 ^K221L^* gene | This study |
| 9 | pINA1269- aroG^G146N^ | pINA1269 plasmid harbors *aroG^G146N^* gene | This study |
| 10 | pINA1312-CAR-SFP | pINA1312 plasmid harbors *CAR* and *sfp* genes | This study |
| 11 | pINA1312-CAR-SFP-AS | pINA1312 plasmid harbors *CAR*, *sfp*, and *AS* genes | This study |
| 12 | pINA1312-UbiC-CAR-SFP | pINA1312 plasmid harbors *ubiC*, *CAR*, and *sfp* genes | This study |
| 13 | pINA1269-UbiC-AS | pINA1269 plasmid harbors *ubiC* and *AS* genes | This study |
| 14 | pINA1269-UbiC-DHS1-AS | pINA1269 plasmid harbors *ubiC*, *DHS1*, and *AS* genes | This study |
| 15 | pINA1269-UbiC-DHS2-AS | pINA1269 plasmid harbors *ubiC*, *DHS2*, and *AS* genes | This study |
| 16 | pINA1269-UbiC-ylARO4^K221L^-AS | pINA1269 plasmid harbors *ubiC*, *ylARO4^K221L^*, and *AS* genes | This study |
| 17 | pINA1269-UbiC-aroG^G146N^-AS | pINA1269 plasmid harbors *ubiC*, *aroG^G146N^*, and *AS* genes | This study |
| 18 | pINA1269-UbiC-DHS1-DHS2-AS | pINA1269 plasmid harbors *ubiC*, *DHS1*, *DHS2*, and *AS* genes | This study |
| 19 | pINA1269-UbiC-DHS1-ylARO4^K221L^-AS | pINA1269 plasmid harbors *ubiC*, *DHS1*, *ylARO4^K221L^*, and *AS* genes | This study |
| 20 | pINA1269-UbiC-DHS1-aroG^G146N^-AS | pINA1269 plasmid harbors *ubiC*, *DHS1*, *aroG^G146N^*, and *AS* genes | This study |
| 21 | pINA1269-UbiC-DHS2-ylARO4^K221L^-AS | pINA1269 plasmid harbors *ubiC*, *DHS2*, *ylARO4^K221L^*, and *AS* genes | This study |
| 22 | pINA1269-UbiC-DHS2-aroG^G146N^-AS | pINA1269 plasmid harbors *ubiC*, *DHS1*, *aroG^G146N^*, and *AS* genes | This study |
| 23 | pINA1269-UbiC-ylARO4^K221L^-aroG^G146N^-AS | pINA1269 plasmid harbors *ubiC*, *ylARO4^K221L^*, *aroG^G146N^*, and *AS* genes | This study |
| 24 | pINA1269-UbiC-DHS1-DHS2-ylARO4^K221L^-AS | pINA1269 plasmid harbors *ubiC*, *DHS1*, *DHS2*, *ylARO4^K221L^*, and *AS* genes | This study |
| 25 | pINA1269-UbiC-DHS1-DHS2-aroG^G146N^-AS | pINA1269 plasmid harbors *ubiC*, *DHS1*, *DHS2*, *aroG^G146N^*, and *AS* genes | This study |
| 26 | pINA1269-UbiC-DHS1-ylARO4^K221L^-aroG^G146N^-AS | pINA1269 plasmid harbors *ubiC*, *DHS1*, *ylARO4^K221L^*, *aroG^G146N^*, and *AS* genes | This study |
| 27 | pINA1269-UbiC-DHS2-ylARO4^K221L^-aroG^G146N^-AS | pINA1269 plasmid harbors *ubiC*, *DHS2*, *ylARO4^K221L^*, *aroG^G146N^*, and *AS* genes | This study |
| 28 | pINA1269-UbiC-DHS1-DHS2-ylARO4^K221L^-aroG^G146N^-AS | pINA1269 plasmid harbors *ubiC*, *DHS1*, *DHS2*, *ylARO4^K221L^*, *aroG^G146N^*, and *AS* genes | This study |
| 29 | pYlCas9 | pRRQ2 plasmid harbors *Cas9* genes | This study |
| 30 | pYlCas9-ΔylTRP1 | pRRQ2 plasmid harbors *Cas9* genes and gRNA（ΔylTRP1） | This study |
| 31 | pYlCas9-ΔylARO7 | pRRQ2 plasmid harbors *Cas9* genes and gRNA（ΔylARO7） | This study |
| 32 | pYlCas9-ΔylARO10 | pRRQ2 plasmid harbors *Cas9* genes and gRNA（ΔylARO10） | This study |
| 33 | pYlCas9-ΔylPAR4 | pRRQ2 plasmid harbors *Cas9* genes and gRNA（ΔylPAR4） | This study |
| 34 | pYlCas9-ΔylPHA2 | pRRQ2 plasmid harbors *Cas9* genes and gRNA（ΔylPHA2） | This study |
| 35 | pYlCas9-ΔylTYR1 | pRRQ2 plasmid harbors *Cas9* genes and gRNA（ΔylTYR1） | This study |

**Table S2.** Primers used in this study

| Name | | Sequence |
| --- | --- | --- |
| 1 | AS-F | ATGGAACACACCCCTCACATTGCCATG |
|  | AS-R | TTAGGTAGAAGAGATCTTGTTCTCCCACTTGC |
|  | 1312-AS-F | ACAAGATCTCTTCTACCTAAGGTACCTCCATGGCCTGTCCCCAC |
|  | 1312-AS-R | CACACATACAACCACACACATCCACGTGATGGAACACACCCCTCACAT |
| 2 | 1269-AS-F | ACAAAATTAGCAGCACCTAAGGTACCTCCATGGCCTGTCCCCAC |
|  | 1269-AS-R | ATATGCGGTGTATGTTCCATCACGTGGATGTGTGTGGTTGTATGTG |
| 3 | CAR-F | TTTTGCAGTACTAACCGCAGACCGAGACTATCTCTACCGCCGCCGT |
|  | CAR-R | TTAGACCAGACCCAGCAGCTGGATGTC |
|  | 1312-CAR-F | AGCTGCTGGGTCTGGTCTAACACGTGGGAACCCGAAACTAAGGATC |
|  | 1312-CAR-R | GCAGCCCAAGCTAGCTTATCGATACGCGT |
| 4 | SFP-F | CACATACAACCACACACATCATGAAGATCTACGGTATTTACATGGACCGAC |
|  | SFP-R | TTACAGCAGCTCCTCGTAGGACACC |
|  | 1312-SFP-F | CCTACGAGGAGCTGCTGTAACACGTGGGAACCCGAAACTAAGGATC |
|  | 1312-SFP-R | GATGTGTGTGGTTGTATGTGTGATGTGG |
| 5 | UbiC-F | TTTTGCAGTACTAACCGCAGTCTCACCCCGCTCTCACCCAGC |
|  | UbiC-R | TTAGTACAGGGGAGAAGCAGGCAGGAAC |
|  | 1269-UbiC-F | CTGCTTCTCCCCTGTACTAAAACTACGGAACTTGTGTTGATGTCTTTGCC |
|  | 1269-UbiC-R | ATTCCTTGCGGCGGCGGTGCTC |
| 6 | DHS1-F | CCAGTAGTAGGTTGAGGCCG |
|  | DHS1-R | ACACGGGCATCTCACTTGCATATGTATG |
|  | 1269-DHS1-F | CATACATATGCAAGTGAGATGCCCGTGTCCGAATTCTCATGTTTGACAGC |
|  | 1269-DHS1-R | CGGCCTCAACCTACTACTGGGCTGCTTCCTAATGCAGGAG |
| 7 | DHS2-F | GGAACCCGAAACTAAGGATCATGCCACCCAAAGTCGTTATTACCGATC |
|  | DHS2-R | CAACACAAGTTCCGTAGTTGCTACTTCTGGGCTCTCATGCTGTCTGCA |
|  | 1269-DHS2-F | CAACTACGGAACTTGTGTTGATGTCTTTGC |
|  | 1269-DHS2-R | GATCCTTAGTTTCGGGTTCCCACGTGGATG |
| 8 | ylARO4 ^K221L^ -F | TAGTAGGTTGAGGCCGTTGAGCACCGCCGCCGCAAGGAAT |
|  | ylARO4 ^K221L^ -R | GCTTATCATCGATGATAAGCTGTCAAACATGAGAATTCGG |
|  | 1269-ylARO4 ^K221L^ -F | GCTTATCATCGATGATAAGC |
|  | 1269-ylARO4 ^K221L^ -R | TCAACGGCCTCAACCTACTA |
| 9 | aroG^G146N^-F | GTAGTAGGTTGAGGCCGTTGAGCACCGCCGCCGCAAGGAAT |
|  | aroG^G146N^ -R | GCTTATCATCGATGATAAGCTGTCAAACATGAGAATTCGG |
|  | 1269- aroG^G146N^ -F | GCTTATCATCGATGATAAGC |
|  | 1269- aroG^G146N^ -R | TCAACGGCCTCAACCTACTAC |
| 10 | CAR-F | GCTAGCTTATCGATACGCGTAGAGACCGGGTTGGCGGCGC |
|  | CAR-R | GGAGCTGCATGTGTCAGAGGCCGGGCATCTCACTTGCGTATG |
|  | V4-F | CCTCTGACACATGCAGCTCC |
|  | V4-R | ACGCGTATCGATAAGCTAGC |
| 11 | AS-F | CCCGTGTCCCAAGCTAGCTTATCCCAAGCTAGCTTATCGATACGCGTG |
|  | AS-R | CCAGAGCGAGTGTTACACATGGAATTCCATCTCACTTGCGTATGTATGG |
|  | V10-F | GAATTCCATGTGTAACACTCGCTCTGG |
|  | V10-R | GGATAAGCTAGCTTGGGACACGGG |
| 12 | UbiC-F | GCTAGCTTATCGATACGCGTAGAGACCGGGTTGGCGGCGC |
|  | UbiC-R | ACACGGGCATCTCACTTGC |
|  | V10-F | GCAAGTGAGATGCCCGTGTAGAGACCGGGTTGGCGGCGC |
|  | V10-R | ACGCGTATCGATAAGCTAGC |
| 13 | AS-F | GCACCGCCGCCGCAAGGAAT |
|  | AS-R | GCGCCGCCAACCCGGTCTCTACACGGGCATCTCACTTGCA |
|  | V5-F | AGAGACCGGGTTGGCGGCGC |
|  | V5-R | ATTCCTTGCGGCGGCGGTGCTCAACACTAGTGGATCTGCTG |
| 14 | DHS1-F | TGCAAGTGAGATGCCCGTGTGCACCGCCGCCGCAAGGAAT |
|  | DHS1-R | GCGCCGCCAACCCGGTCTCTACACGGGCATCTCACTTGCATATG |
|  | V13-F | AGAGACCGGGTTGGCGGCGC |
|  | V13-R | ACACGGGCATCTCACTTGCA |
| 15 | DHS2-F | CCCTTATGCGACTCCTGTTGAGCACCGCCGCCGCAAGGAAT |
|  | DHS2-R | GTTTGACAGCTTATCATCGATGATAAGCTGTCAAACATGAGAATTCGG |
|  | V13-F | CCGAATTCTCATGTTTGACAGCTTATCATCGATGATAAGCTGTCAAAC |
|  | V13-R | CAACAGGAGTCGCATAAGGG |
| 16 | ylARO4^K221L^-F | CTTATGCGACTCCTGCATTAGGAAGCAGCCCAGTAGTAGGTTGAG |
|  | ylARO4^K221L^-R | TTAGTTCTTGTTTCGTCGCTCCTTGACGGCGTTGGC |
|  | V13-F | AGCGACGAAACAAGAACTAACAACTACGGAACTTGTGTTGATGTCTTTGCCCCCGG |
|  | V13-R | CCTAATGCAGGAGTCGCATAAG |
| 17 | aroG^G146N^-F | GGAACCCGAAACTAAGGATCATGAATTATCAGAACGACGATTTACGCATCAAAGAAATC |
|  | aroG^G146N^-R | TTACCCGCGACGCGCTTTTACTGCATTC |
|  | V13-F | TAAAAGCGCGTCGCGGGTAACAACTACGGAACTTGTGTTGATGTCTTTGCCCCCGG |
|  | V13-R | GATCCTTAGTTTCGGGTTCCCACGTGGATG |
| 18 | DHS2-F | TGCAAGTGAGATGCCCGTGTCCGAATTCTCATGTTTGACA |
|  | DHS2-R | GAGACACCTCAGCATGCACCATTCCTTGCGGCGGCGGTGC |
|  | V14-F | AACAGTGTACGCAGTACTAT |
|  | V14-R | ACGCAACTAACATGAATGAA |
| 19 | ylARO4^K221L^-F | GCCGCCTAGATGACAAATTC |
|  | ylARO4^K221L^-R | ATTCCTTGCGGCGGCGGTGCTCAACGGCCTCAACCTACTAC |
|  | V14-F | GCACCGCCGCCGCAAGGAAT |
|  | V14-R | GAATTTGTCATCTAGGCGGCCTGGCCGTCTTCTCCGGGGC |
| 20 | aroG^G146N^-F | GACGCAGTAGGATGTCCTGC |
|  | aroG^G146N^-R | CATCTCACTTGCGTATGTATG |
|  | V14-F | CATACATACGCAAGTGAGATGGGTGCCTAATGAGTGAGCTAAC |
|  | V14-R | GCAGGACATCCTACTGCGTCTGCTTCCTAATGCAGGAGTCGC |
| 21 | DHS2-F | GCACCGCCGCCGCAAGGAAT |
|  | DHS2-R | ACACGGGCATCTCACTTGCA |
|  | V16-F | TGCAAGTGAGATGCCCGTGTCCGAATTCTCATGTTTGACAG |
|  | V16-R | ATTCCTTGCGGCGGCGGTGCTCAACGGCCTCAACCTACTAC |
| 22 | DHS2-F | GCACCGCCGCCGCAAGGAAT |
|  | DHS2-R | ACACGGGCATCTCACTTGCA |
|  | V17-F | TGCAAGTGAGATGCCCGTGTCCGAATTCTCATGTTTGACAG |
|  | V17-R | ATTCCTTGCGGCGGCGGTGCACACGGGCATCTCACTTGCATATG |
| 23 | ylARO4^K221L^-F | CTTATGCGACTCCTGCATTAGGAAGCAGCCCAGTAGTAGGTTGAG |
|  | ylARO4^K221L^-R | ATTCCTTGCGGCGGCGGTGCTGTCAAACATGAGAATTCGG |
|  | V17-F | GCACCGCCGCCGCAAGGAAT |
|  | V17-R | CCTAATGCAGGAGTCGCATAAG |
| 24 | ylARO4^K221L^-F | GACTCCTGCATTAGGAAGCAAACAGTGTACGCAGTACTATAGAGGAACAATTGC |
|  | ylARO4^K221L^-R | ACGCAACTAACATGAATGAATACGATATACATCAAAG |
|  | V18-F | TTCATTCATGTTAGTTGCGTGTTGAGCACCGCCGCCGCAAGGAAT |
|  | V18-R | TGCTTCCTAATGCAGGAGTCGCATAAGGGAG |
| 25 | aroG^G146N^-F | GACTCCTGCATTAGGAAGCAGACGCAGTAGGATGTCCTGCACGGGTCTTTTTG |
|  | aroG^G146N^-R | CAATTAATGTGAGTTAGCTCACTCATTAGGCACC |
|  | V18-F | GAGCTAACTCACATTAATTGGTTGAGCACCGCCGCCGCAAGGAAT |
|  | V18-R | TGCTTCCTAATGCAGGAGTCGCATAAGGGAG |
| 26 | ARO4 ^K221L^ -F | GAGCTAACTCACATTAATTGAACAGTGTACGCAGTACTATAGAGGAACAATTGC |
|  | ARO4 ^K221L^ -R | ACGCAACTAACATGAATGAATACGATATACATCAAAG |
|  | V20-F | TTCATTCATGTTAGTTGCGTGCCCAGTAGTAGGTTGAGGCCGTTG |
|  | V20-R | CAATTAATGTGAGTTAGCTCACTCATTAGGCACC |
| 27 | ARO4 ^K221L^ -F | GAGCTAACTCACATTAATTGAACAGTGTACGCAGTACTATAGAGGAACAATTGC |
|  | ARO4 ^K221L^ -R | ACGCAACTAACATGAATGAATACGATATACATCAAAG |
|  | V22-F | TTCATTCATGTTAGTTGCGTGCCCAGTAGTAGGTTGAGGCCGTTG |
|  | V22-R | CAATTAATGTGAGTTAGCTCACTCATTAGGCACC |
| 28 | DHS2-F | TTCATTCATGTTAGTTGCGTGTTGAGCACCGCCGCCGCAAGGAAT |
|  | DHS2-R | GAGACACGGCTCCGCCAGAGACACGGGCATCTCACTTGCATATGTATGG |
|  | V26-F | CTCTGGCGGAGCCGTGTCTCGCCCAGTAGTAGGTTGAGGCCGTTG |
|  | V26-R | ACGCAACTAACATGAATGAATACGATATACATCAAAG |
| 29 | Cas9-F | ATGGACAGGTAGTAGGAGGCAAG |
|  | Cas9-R | TGCTGCGGTAAAGCTCATCAG |
|  | gRNA-F | CCCCAGTTGCAAAAGTTGACAC |
|  | gRNA-R | GCCTCCTACTACCTGTCCATAAAAAAAAGCACCGACTCGGTGCCAC |
|  | V-F | CTGATGAGCTTTACCGCAGCA |
|  | V-R | GTGTCAACTTTTGCAACTGGGGCACACTCCTTTGACATAACG |
| 30 | SgTRP1-F | TGTTTGTCAAAAACGCCAGGGTTTTAGAGCTAGAAATAGCAAGTTAAAATAAGGCTAG |
|  | SgTRP1-R | CCTGGCGTTTTTGACAAACAACGTCAACCTGCGCCGAC |
| TRP1 homology arm | TRP1up-F | CTCGCTCATTCCAGCGGTTG |
|  | TRP1up-R | CACATCCGCCGCTGTTTACGTCCATGATGACAGTGGCGAG |
|  | TRP1down-F | CTCGCCACTGTCATCATGGACGTAAACAGCGGCGGATGTG |
|  | TRP1down-R | CTGGACGACGTACGTGTGTC |
| 31 | SgARO7-F | GTAGATTCGCTTGACAACCTGTTTTAGAGCTAGAAATAGCAAGTTAAAATAAGGC |
|  | SgARO7-R | AGGTTGTCAAGCGAATCTACGACGAGCTTACTCGTTTCGTCC |
| ARO7 homology arm | ARO7up-F | ACAAGCTGTTCAACGGACCC |
|  | ARO7up-R | CTGCCCAAATTCGCCGGAAACGTGTGTTTTTTTAGCGAAGCGAG |
|  | ARO7down-F | TTTCCGGCGAATTTGGGCAG |
|  | ARO7down-R | GTTTGGCGGGCGCTTAGAC |
| 32 | SgARO10-F | GCTGCCATGTCATATCCACGGTTTTAGAGCTAGAAATAGCAAGTTAAAATAAGGCTAG |
|  | SgARO10-R | CGTGGATATGACATGGCAGCACGTCAACCTGCGCCGAC |
| ARO10 homology arm | ARO10up-F | GGATGCCTAAGGAGCCCG |
|  | ARO10up-R | ATGATGAGATTGAGCGGAGCG |
|  | ARO10down-F | CGCTCCGCTCAATCTCATCATGTTGATAGTCAAGTCACTGGAGAGATGG |
|  | ARO10down-R | GTATGTACGATAGTCGTCTGATCAGGTG |
| 33 | SgPAR4-F | CTCGTGCCAGTTCATCATGGGTTTTAGAGCTAGAAATAGCAAGTTAAAATAAGGC |
|  | SgPAR4-R | CCATGATGAACTGGCACGAGACGTCAACCTGCGCCGAC |
| PAR4 homology arm | PAR4up-F | GCTGACCGAATCCGAAAAGAGC |
|  | PAR4up-R | CAAACACAACAACGTTCGAGAGCGAAGCC |
|  | PAR4down-F | CTCGAACGTTGTTGTGTTTGTGTGTTGGATGTGTG |
|  | PAR4down-R | CCAGCACTTCTAACATACAGTATTGTAGC |
| 34 | SgPHA2-F | CTCGGGAGAAGTCAAGCTGGGTTTTAGAGCTAGAAATAGCAAGTTAAAATAAGGCTAG |
|  | SgPHA2-R | CCAGCTTGACTTCTCCCGAGACGTCAACCTGCGCCGAC |
| PHA2 homology arm | PHA2up-F | CACTCTGGCCGAGTTCAAGC |
|  | PHA2up-R | GGTGATGTGTAATGTGTGTGATCAAGTG |
|  | PHA2down-F | CACTTGATCACACACATTACACATCACCACGGTTCAGCGTTTCTGTCTAGG |
|  | PHA2down-R | TGAGAGAATACCTTGATTCTGGCCACC |
| 35 | SgTYR1-F | TGCACTGTTCGAGCAGTCCTGTTTTAGAGCTAGAAATAGCAAGTTAAAATAAGGCTAG |
|  | SgTYR1-R | AGGACTGCTCGAACAGTGCAACGTCAACCTGCGCCGAC |
| TYR1 homology arm | TYR1up-F | CCTCCGAAGAGGCTCTCAAAATG |
|  | TYR1up-R | GACACACTTGCAGGTCTAAAAGTTCC |
|  | TYR1down-F | GGAACTTTTAGACCTGCAAGTGTGTCGTTGTAGAGCGTGGCGAAAG |
|  | TYR1down-R | GGACAGAGTGTCCAACAAGCC |
| gRNA used in this study | gRNA-TRP1 | TGTTTGTCAAAAACGCCAGG |
|  | gRNA-ARO7 | GTAGATTCGCTTGACAACCT |
|  | gRNA-ARO10 | GCTGCCATGTCATATCCACG |
|  | gRNA-PAR4 | CTCGTGCCAGTTCATCATGG |
|  | gRNA-PHA2 | CTCGGGAGAAGTCAAGCTGG |
|  | gRNA-TYR1 | TGCACTGTTCGAGCAGTCCT |
|  | gRNA-MHY1 | GGACGCCGTTTCCATCTCAC |
|  | gRNA-CLA4 | TTCACACATAAGGTGCACGT |

The numbers in table S2 correspond in table S1.

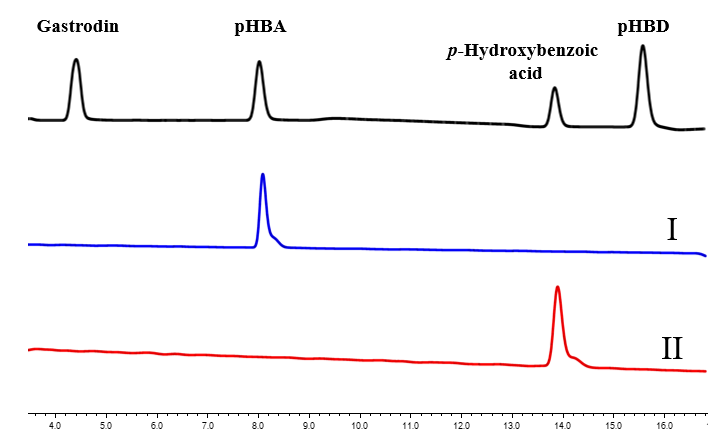

**Figure S1.** The supernatant from strain Po1fΔ*Ku*70 feeding pHBA (I) and *p*-hydroxybenzoic acid (II) were subjected to HPLC analysis, respectively.

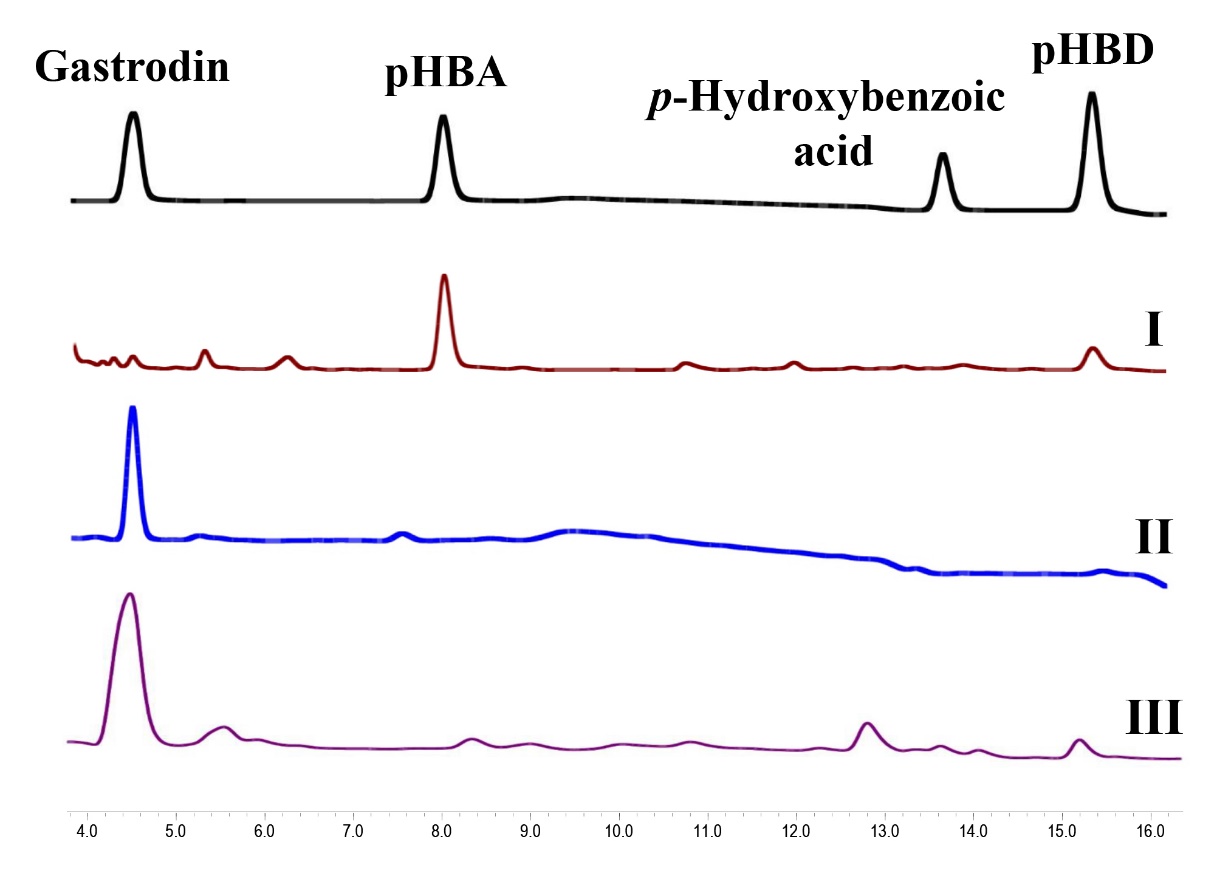

**Figure S2.** The supernatant from the recombinant strains SC03 (I), Gd01 (II) and Gd07 (II) were subjected to HPLC analysis, respectively.

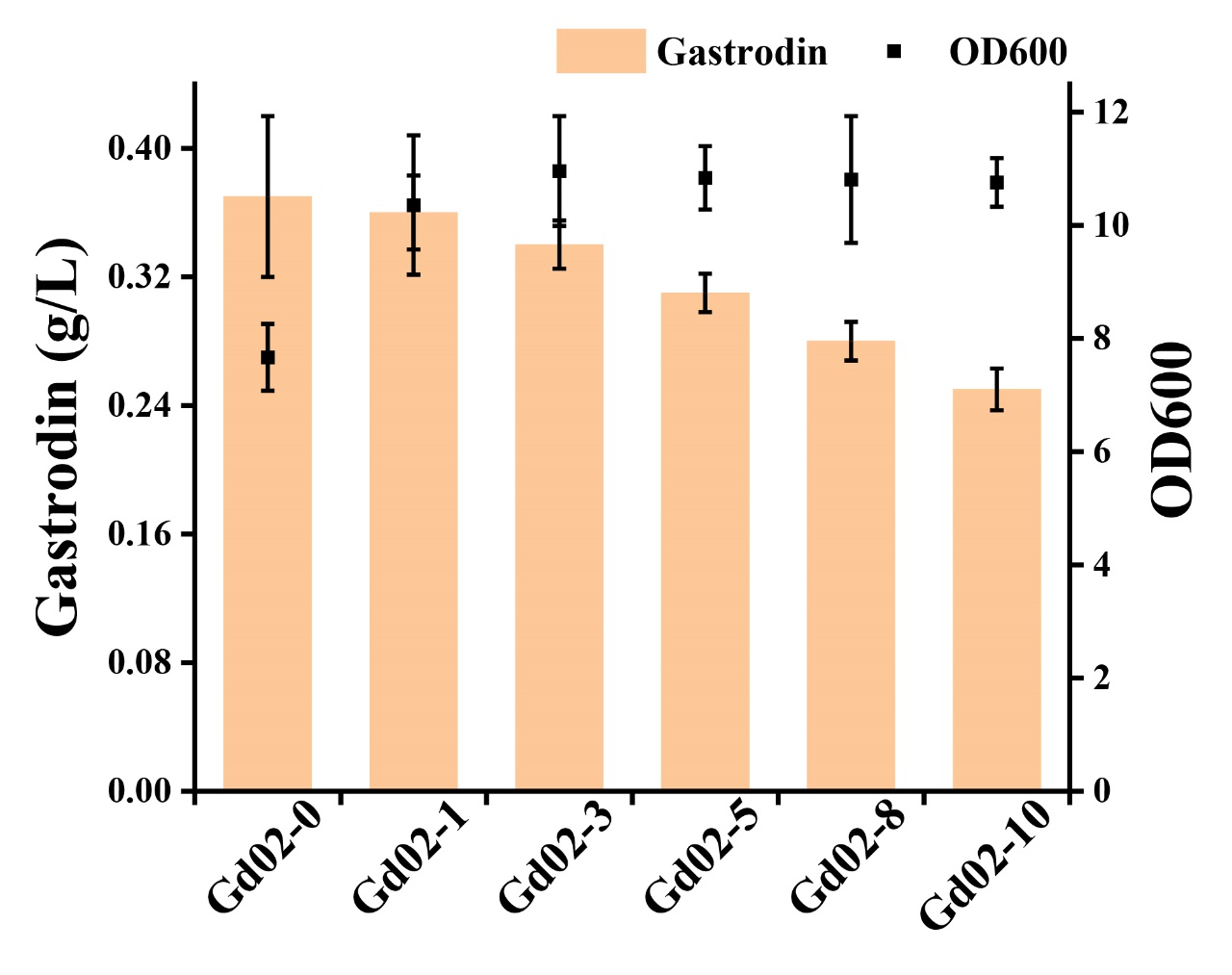

**Figure S3.** The effects of different tryptophan concentrations on gastrodin and biomass production of strain Gd03. 0, 1, 3, 5, 8, 10 represent 0 g/L, 1 g/L, 3 g/L, 5 g/L, 8 g/L, and 10 g/L of tryptophan adding into culture.

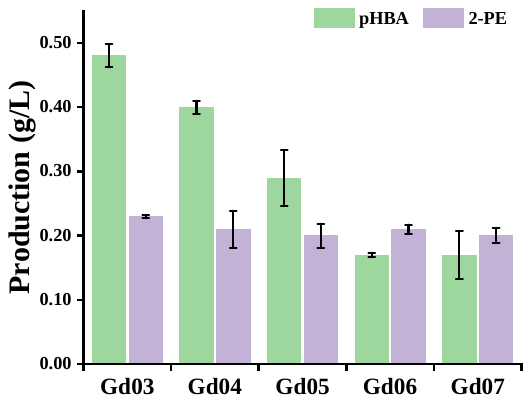

**Figure S4.** The productions of pHBA and 2-PE produced by different strains.

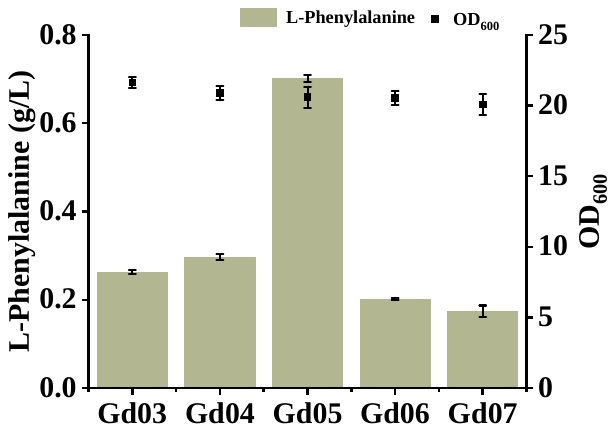
**Figure S5.** The productions of phenylalanine produced by different strains.

pINA1312

AAGCTAGCTTATCGATACGCGTGCATGCTGAGGTGTCTCACAAGTGCCGTGCAGTCCCGCCCCCACTTGCTTCTCTTTGTGTGTAGTGTACGTACATTATCGAGACCGTTGTTCCCGCCCACCTCGATCCGGCATGCTGAGGTGTCTCACAAGTGCCGTGCAGTCCCGCCCCCACTTGCTTCTCTTTGTGTGTAGTGTACGTACATTATCGAGACCGTTGTTCCCGCCCACCTCGATCCGGCATGCTGAGGTGTCTCACAAGTGCCGTGCAGTCCCGCCCCCACTTGCTTCTCTTTGTGTGTAGTGTACGTACATTATCGAGACCGTTGTTCCCGCCCACCTCGATCCGGCATGCTGAGGTGTCTCACAAGTGCCGTGCAGTCCCGCCCCCACTTGCTTCTCTTTGTGTGTAGTGTACGTACATTATCGAGACCGTTGTTCCCGCCCACCTCGATCCGGCATGCACTGATCACGGGCAAAAGTGCGTATATATACAAGAGCGTTTGCCAGCCACAGATTTTCACTCCACACACCACATCACACATACAACCACACACATCCACGTGGGAACCCGAAACTAAGGATCCGGTACCTCCATGGCCTGTCCCCACGTTGCCGGTCTTGCCTCCTACTACCTGTCCATCAATGACGAGGTTCTCACCCCTGCCCAGGTCGAGGCTCTTATTACTGAGTCCAACACCGGTGTTCTTCCCACCACCAACCTCAAGGGCTCTCCCAACGCTGTTGCCTACAACGGTGTTGGCATTTAGGCAATTAACAGATAGTTTGCCGGTGATAATTCTCTTAACCTCCCACACTCCTTTGACATAACGATTTATGTAACGAAACTGAAATTTGACCAGATATTGTTGTAAATAGAAAATCTGGCTTGTAGGTGGCAAAATCCCGTCTTTGTTCATCAATTCCCTCTGTGACTACTCGTCATCCCTTTATGTTCGACTGTCGTATTTTTATTTTCCATACATACGCAAGTGAGATGCCCGTGTCCGAATTCCATGTGTAACACTCGCTCTGGAGAGTTAGTCATCCGACAGGGTAACTCTAATCTCCCAACACCTTATTAACTCTGCGTAACTGTAACTCTTCTTGCCACGTCGATCTTACTCAATTTTCCTGCTCATCATCTGCTGGATTGTTGTCTATCGTCTGGCTCTAATACATTTATTGTTTATTGCCCAAACAACTTTCATTGCACGTAAGTGAATTGTTTTATAACAGCGTTCGCCAATTGCTGCGCCATCGTCGTCCGGCTGTCCTACCGTTAGGGTAGTGTGTCTCACACTACCGAGGTTACTAGAGTTGGGAAAGCGATACTGCCTCGGACACACCACCTGGTCTTACGACTGCAGAGAGAATCGGCGTTACCTCCTCACAAAGCCCTCAGTGCGGCCGCCCGGGGTGGGCGAAGAACTCCAGCATGAGATCCCCGCGCTGGAGGATCATCCAGCCGGCGTCCCGGAAAACGATTCCGAAGCCCAACCTTTCATAGAAGGCGGCGGTGGAATCGAAATCTCGTGATGGCAGGTTGGGCGTCGCTTGGTCGGTCATTTCGAACCCCAGAGTCCCGCTCAGAAGAACTCGTCAAGAAGGCGATAGAAGGCGATGCGCTGCGAATCGGGAGCGGCGATACCGTAAAGCACGAGGAAGCGGTCAGCCCATTCGCCGCCAAGCTCTTCAGCAATATCACGGGTAGCCAACGCTATGTCCTGATAGCGGTCCGCCACACCCAGCCGGCCACAGTCGATGAATCCAGAAAAGCGGCCATTTTCCACCATGATATTCGGCAAGCAGGCATCGCCATGGGTCACGACGAGATCCTCGCCGTCGGGCATGCGCGCCTTGAGCCTGGCGAACAGTTCGGCTGGCGCGAGCCCCTGATGCTCTTCGTCCAGATCATCCTGATCGACAAGACCGGCTTCCATCCGAGTACGTGCTCGCTCGATGCGATGTTTCGCTTGGTGGTCGAATGGGCAGGTAGCCGGATCAAGCGTATGCAGCCGCCGCATTGCATCAGCCATGATGGATACTTTCTCGGCAGGAGCAAGGTGAGATGACAGGAGATCCTGCCCCGGCACTTCGCCCAATAGCAGCCAGTCCCTTCCCGCTTCAGTGACAACGTCGAGCACAGCTGCGCAAGGAACGCCCGTCGTGGCCAGCCACGATAGCCGCGCTGCCTCGTCCTGCAGTTCATTCAGGGCACCGGACAGGTCGGTCTTGACAAAAAGAACCGGGCGCCCCTGCGCTGACAGCCGGAACACGGCGGCATCAGAGCAGCCGATTGTCTGTTGTGCCCAGTCATAGCCGAATAGCCTCTCCACCCAAGCGGCCGGAGAACCTGCGTGCAATCCATCTTGTTCAATCATGCGAAACGATCCTCATCCTGTCTCTTGATCAGATCTTGATCCCCTGCGCCATCAGATCCTTGGCGGCAAGAAAGCCATCCAGTTTACTTTGCAGGGCTTCCCAACCTTACCAGAGGGCGCCCCAGCTGGCAATTCCGGTTCGCTTGCTGTCCATAAAACCGCCCAGTCTAGCTATCGCCATGTAAGCCCACTGCAAGCTACCTGCTTTCTCTTTGCGCTTGCGTTTTCCCTTGTCCAGATAGCCCAGTAGCTGACATTCATCCGGGGTCAGCACCGTTTCTGCGGACTGGCTTTCTACGTGTTCCGCTTCCTTTAGCAGCCCTTGCGCCCTGAGTGCTTGCGGCAGCGTGAAGCTAGCTTATGCGGTGTGAAATACCGCACAGATGCGTAAGGAGAAAATACCGCATCAGGCGCTCTTCCGCTTCCTCGCTCACTGACTCGCTGCGCTCGGTCGTTCGGCTGCGGCGAGCGGTATCAGCTCACTCAAAGGCGGTAATACGGTTATCCACAGAATCAGGGGATAACGCAGGAAAGAACATGTGAGCAAAAGGCCAGCAAAAGGCCAGGAACCGTAAAAAGGCCGCGTTGCTGGCGTTTTTCCATAGGCTCCGCCCCCCTGACGAGCATCACAAAAATCGACGCTCAAGTCAGAGGTGGCGAAACCCGACAGGACTATAAAGATACCAGGCGTTTCCCCCTGGAAGCTCCCTCGTGCGCTCTCCTGTTCCGACCCTGCCGCTTACCGGATACCTGTCCGCCTTTCTCCCTTCGGGAAGCGTGGCGCTTTCTCATAGCTCACGCTGTAGGTATCTCAGTTCGGTGTAGGTCGTTCGCTCCAAGCTGGGCTGTGTGCACGAACCCCCCGTTCAGCCCGACCGCTGCGCCTTATCCGGTAACTATCGTCTTGAGTCCAACCCGGTAAGACACGACTTATCGCCACTGGCAGCAGCCACTGGTAACAGGATTAGCAGAGCGAGGTATGTAGGCGGTGCTACAGAGTTCTTGAAGTGGTGGCCTAACTACGGCTACACTAGAAGGACAGTATTTGGTATCTGCGCTCTGCTGAAGCCAGTTACCTTCGGAAAAAGAGTTGGTAGCTCTTGATCCGGCAAACAAACCACCGCTGGTAGCGGCGGTTTTTTGTTTGCAAGCAGCAGATTACGCGCAGAAAAAAAGGATCTCAAGAAGATCCTTTGATCTTTTCTTACTGAACGGTGATCCCCACCGGAATTGCGGCCGCTGTCGGGAACCGCGTTCAGGTGGAACAGGACACCTCCCTTGCACTTCTTGGTATATCAGTATAGGCTGATGTATTCATAGTGGGGTTTTTCATAATAAATTTACTAACGGCAGGCAACATTCACTCGGCTTAAACGCAAAACGGACCGTCTTGATATCTTCTGACGCATTGACCACCGAGAAATAGTGTTAGTTACCGGGTGAGTTATTGTTCTTCTACACAGGCGACGCCCATCGTCTAGAGTTGATGTACTAACTCAGATTTCACTACCTACCCTATCCCTGGTACGCACAAAGCACTTTGCTAGATAGAGTCGACAAAGGCCTGTTTCTCGGTGTACAGAGCTTGGTCCTCCTTGAAGTTGCGACACATGTCTTGATAGTATCTTGGCTTCTCTCTCTTGAGCTTTTCCATAACAAGTTCTTCTGCCTCCAGGAAGTCCATGGGTGGTTTGATCATGGTTTTGGTGTAGTGGTAGTGCAGTGGTGGTATTGTGACTGGGGATGTAGTTGAGAATAAGTCATACACAAGTCAGCTTTCTTCGAGCCTCATATAAGTATAAGTAGTTCAACGTATTAGCACTGTACCCAGCATCTCCGTATCGAGAAACACAACAACATGCCCCATTGGACAGACCATGCGGATACACAGGTTGTGCAGTACCATACATACTCGATCAGACAGGTCGTCTGACCATCATACAAGCTGAACAGCGCTCCATACTTGCACGCTCTCTATATACACAGTTAAATTACATATCCATAGTCTAACCTCTAACAGTTAATCTTCTGGTAAGCCTCCCAGCCAGCCTTCTGGTATCGCTTGGCCTCCTCAATAGGATCTCGGTTCTGGCCGTACAGACCTCGGCCGACAATTATGATATCCGTTCCGGTAGACATGACATCCTCAACAGTTCGGTACTGCTGTCCGAGAGCGTCTCCCTTGTCGTCAAGACCCACCCCGGGGGTCAGAATAAGCCAGTCCTCAGAGTCGCCCTTAGGTCGGTTCTGGGCAATGAAGCCAACCACAAACTCGGGGTCGGATCGGGCAAGCTCAATGGTCTGCTTGGAGTACTCGCCAGTGGCCAGAGAGCCCTTGCAAGACAGCTCGGCCAGCATGAGCAGACCTCTGGCCAGCTTCTCGTTGGGAGAGGGGACTAGGAACTCCTTGTACTGGGAGTTCTCGTAGTCAGAGACGTCCTCCTTCTTCTGTTCAGAGACAGTTTCCTCGGCACCAGCTCGCAGGCCAGCAATGATTCCGGTTCCGGGTACACCGTGGGCGTTGGTGATATCGGACCACTCGGCGATTCGGTGACACCGGTACTGGTGCTTGACAGTGTTGCCAATATCTGCGAACTTTCTGTCCTCGAACAGGAAGAAACCGTGCTTAAGAGCAAGTTCCTTGAGGGGGAGCACAGTGCCGGCGTAGGTGAAGTCGTCAATGATGTCGATATGGGTCTTGATCATGCACACATAAGGTCCGACCTTATCGGCAAGCTCAATGAGCTCCTTGGTGGTGGTAACATCCAGAGAAGCACACAGGTTGGTTTTCTTGGCTGCCACGAGCTTGAGCACTCGAGCGGCAAAGGCGGACTTGTGGACGTTAGCTCGAGCTTCGTAGGAGGGCATTTTGGTGGTGAAGAGGAGACTGAAATAAATTTAGTCTGCAGCCC

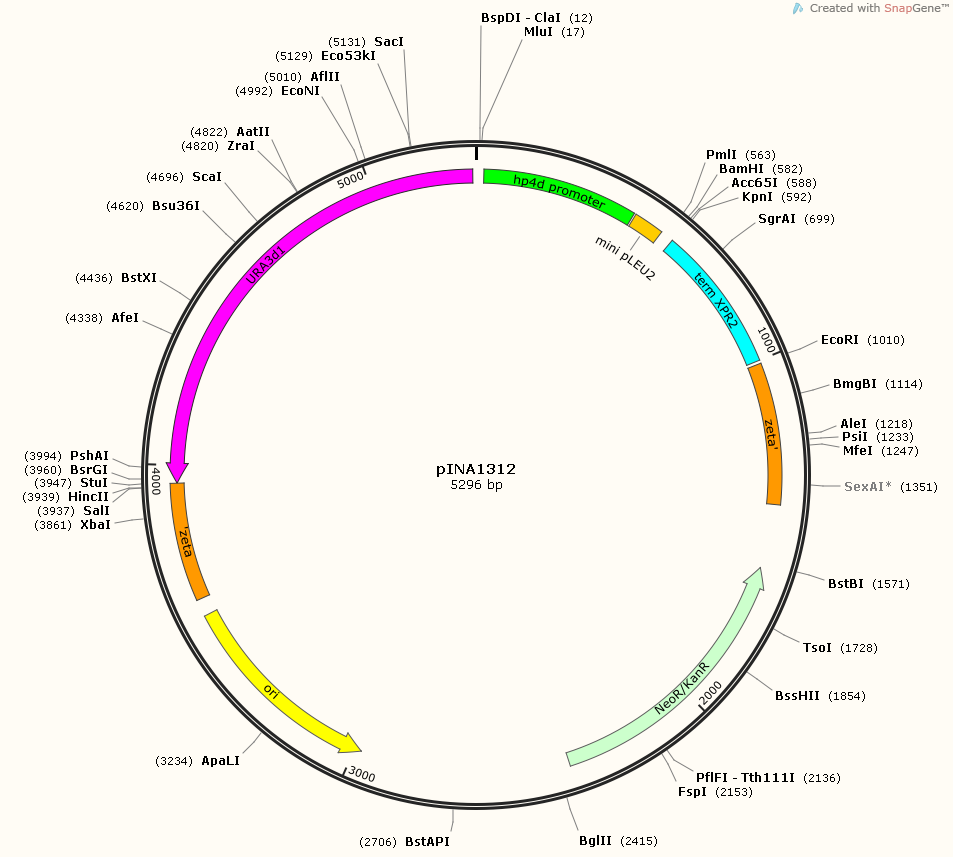

(1) Shang, Y.; Wei, W.; Zhang, P.; Ye, B.-C. Engineering *Yarrowia lipolytica* for enhanced production of arbutin. *J. Agric. Food Chem.* **2020**, *68* (5), 1364-1372.
